# *MIR319C* and its target *JAW-TCP*s repress each other and establish cell proliferation pattern in incipient leaf primordia in *Arabidopsis thaliana*

**DOI:** 10.1101/2023.08.02.551638

**Authors:** Naveen Shankar, Preethi Sunkara, Utpal Nath

## Abstract

The microRNA miR319 and its target JAW-TCP transcription factors regulate leaf morphogenesis in diverse plant species. In young Arabidopsis leaf primordia, *JAW-TCP*s are detected towards the distal region whereas *MIR319C* is expressed at the base. Little is known about how this complementary expression pattern of *MIR319C* and *JAW-TCPs* is generated. Here, we show that *MIR319C* is initially expressed uniformly throughout the incipient primordia and is later abruptly down-regulated at the distal region, with concomitant distal appearance of *JAW-TCP*s, when leaves grow to ∼100 µm long. Loss of *JAW-TCPs* causes distal extension of *MIR319C* expression domain, whereas ectopic TCP activity restricts *MIR319C* more proximally. JAW-TCPs are recruited to and are capable of depositing histone H3K27me3 repressive marks on the *MIR319C* chromatin. *JAW-TCP*s fail to repress *MIR319C* in transgenic seedlings where the TCP-binding *cis*-elements on *MIR319C* are mutated, causing miR319 gain-of-function-like phenotype. Based on these results, we propose a model for growth patterning in leaf primordia wherein *MIR319C* and JAW-TCPs repress each other and divide the uniformly growing primordia into distal differentiation zone and proximal proliferation domain.

**Summary statement:** *JAW-TCPs* transcriptionally repress the microRNA319 encoding gene *MIR319C* to generate their mutually exclusive expression pattern and establish growth polarity during early stages of Arabidopsis leaf primordia.

## Introduction

The morphogenesis of angiosperm leaves from a simple rod-like primordium to maturity involves overlapping phases of growth and differentiation (Rodriguez et al., 2013; Bar and Ori, 2014). In Arabidopsis leaves, growth pattern along the proximodistal axis is established at an early stage (∼150 µm length), where the basal region grows at a higher rate compared to the distal end due to more cell proliferation (Kuchen et al., 2012; Kiezkowski et al., 2019). This polar growth pattern is maintained during the subsequent stages where cells at the base continue to divide, whereas cells towards the distal zone exit division and enter differentiation (Donnelly et al., 1999; Andriankaja et al., 2012). At later stages, the lamina displays more uniform growth as all cells exit proliferation and grow by expansion until maturity (Andriankaja et al., 2012; Kuchen et al., 2012). The establishment and maintenance of this early growth pattern is important for achieving the final shape and size of the leaf (Kuchen et al., 2012).

The gene regulatory network that maintains polar growth in leaves has been described to certain extent (Maugarny Cales and Laufs, 2018, Sarvepalli et al., 2019). Genes such as *GROWTH REGULATING FACTORS* (*GRFs*), *ANGUSTIFOLIA3* (*AN3*) and the cytochrome P450-encoding gene *KLUH* (*KLU*) promote growth by maintaining the proliferative state of the cells in the proximal region (Horiguchi et al., 2005; Anastasiou et al., 2007; Kazama et al., 2010; Ichihashi et al., 2011; Vercruyssen et al., 2014). By contrast, the distally expressed *CINCINNATA-*like *TEOSINTE BRANCHED 1, CYCLOIDEA, PROLIFERATING CELL FACTORS* (*CIN*-*TCP)* genes act as growth repressing factors and initiate leaf maturation (Efroni et al., 2008; Challa et al., 2019). CIN-TCPs activate miR396 expression, which leads to a temporal decline of the level of its cognate *GRF* targets from the distal region (Rodriguez et al., 2010; Schommer et al., 2014). In addition, CIN-TCPs downregulate *AN3* and *GRF5/6* expression independent of miR396 (Rodriguez et al., 2010) to restrict the proliferation potential towards the base of the leaf. Thus, leaf growth polarity is likely to be maintained by the crosstalk between the growth-promoting and the growth-repressing factors. However, how the polar growth pattern in incipient leaf primordia is established is not known.

Several plant microRNA-encoding genes (*MIRNA*) are intergenic and carry *cis*-elements for transcription factor (TF)-mediated regulation (Rogers and Chen, 2013; D’Ario et al., 2017). There are examples where a target TF, whose transcripts are degraded by a microRNA (miRNA), in turn downregulates the transcription of the *MIRNA* to form a double-negative feedback loop (DNFBL; Cai et al., 2013). DNFBLs contribute to bistability where a system exists in one of the two states depending on the directionality of the dominant repression (Tsang et al., 2007). In a typical miRNA-TF DNFBL, the directionality depends almost exclusively on an initial trigger that can transiently increase or decrease either the miRNA or the TF. Due to reciprocal repression, the miRNA or the TF exerts positive feedback on itself and amplifies the response relative to the trigger (Li and Carthew, 2005; Cai et al., 2013). The bistable nature of a miRNA-TF DNFBL makes it an ideal genetic mechanism for generating mutually exclusive expression patterns and regulating growth and development (Fazi et al., 2005; Li and Carthew, 2005; Bracken et al., 2008). For example, during Arabidopsis leaf ontogenesis, miR165/166 and its target TF REVOLUTA suppress each other to generate mutually exclusive expression domains and maintain the abaxial-adaxial polarity (Merelo et al., 2016). Although several miRNA-TF DNFBLs are predicted to exist in Arabidopsis (Megraw et al., 2013), their role in plant development remains unexplored.

The conserved miR319 and its cognate target transcripts encoded by five *CIN-TCP* genes (*TCP2*, *3*, *4*, *10* & *24*, henceforth mentioned as *JAGGED AND WAVY TCPs* or *JAW-TCPs*, Palatnik et al., 2003) act as heterochronic factors in regulating leaf morphogenesis by triggering the transition from division to differentiation in leaf pavement cells (Ori et al., 2007; Efroni et al., 2008; Shleizer-Burko et al., 2011; Challa et al., 2019). In Arabidopsis, ectopic expression of miR319 in the *jaw-D* mutant line, or knocking out of multiple *JAW-TCP*s, extends the cell proliferation zone more distally, yielding larger leaves with excess cells (Palatnik et al., 2003; Efroni et al., 2008; Challa et al., 2019). On the contrary, increased *JAW-TCP* activity reduces cell number and leaf size by advancing the onset of differentiation (Palatnik et al., 2003; Efroni et al., 2008; Schommer et al., 2014; Challa et al., 2019), suggesting the importance of spatiotemporal regulation of *MIR319-JAW-TCP* expression in shaping leaf growth.

In early leaf primordia, the *JAW-TCP* transcripts and their protein products are detected in the distal region where differentiation is initiated (Palatnik et al., 2003; Alvarez et al., 2016). This distal-specific JAW-TCP activity can in part be explained by the expression pattern of the miR319-encoding gene *MIR319C*, which remains restricted to the basal region coinciding with the cell proliferation zone (Nag et al., 2009; Challa et al., 2019). As *MIR319A* and *MIR319B* play minor role in leaf morphogenesis in Arabidopsis (Koyama et al., 2017), leaf growth is perhaps regulated only by miR319c activity (Nag et al., 2009). However, how the basal expression pattern of *MIR319C* is generated in early leaf primordia remains unknown. The most parsimonious proposition of dividing an incipient primordium into a mutually exclusive proximal miR319c-expression domain and a distal JAW-TCP-expression domain would be a DNFBL where they directly suppress each other. In this model the miR319c post-transcriptionally downregulates the level of *JAW-TCPs* (Palatnik et al., 2003), whose protein product would in turn repress *MIR319C* transcription. Here, we have tested this possibility in incipient leaf primordia by biochemical and genetic interventions.

## Results

### Complementary expression pattern of *MIR319C* and *TCP4* is established in early leaf primordia

In a young Arabidopsis leaf primordium, the *MIR319C* promoter (*pMIR319C*) is reported to be active at the basal region, complementary to the *JAW-TCP* transcripts detected towards the distal domain (Palatnik et al., 2003; Nag et al., 2009; Alvarez et al., 2016; Challa et al., 2019). To determine the earliest expression pattern of *MIR319C* in leaves, we examined β-glucuronidase (GUS) activity at the shoot apex of a *pMIR319C::GUS* transgenic line (Nag et al., 2009). In wild type, *MIR319C* promoter activity was detected throughout incipient leaf primordia and was excluded from the shoot meristem (**Fig. 1A**). To determine when this widespread expression becomes restricted to the leaf base, we measured the length of the GUS domain along the proximal-distal axis at various stages of leaf growth and expressed it relative to the total leaf length (**Fig. S1; also see *Methods***). In the 123 young (<120 µm in length) wildtype leaf primordia studied, GUS domain occupied either the entire leaf (34 in total), or up to nearly half of the basal leaf domain (**Fig. 1B, 1F**). The downregulation in *MIR319C* promoter activity towards the distal half appeared to have taken place abruptly within a short span of time when the primordia grew from ∼50 µm to ∼100 µm in length (the faded blue box in **Fig. 1F**), which typically takes nearly 24 hours (Ichihashi et al., 2011; Kierzkowski et al., 2019). Following this transition in *MIR319C* promoter activity pattern, the proportion of GUS domain reduced more slowly to <0.4 when the leaf primordia grew to ∼350 µm long (**Fig. 1F**), and to ∼0.25 in ∼1 mm long leaf (**Fig. S2A**).

**Figure 1.**
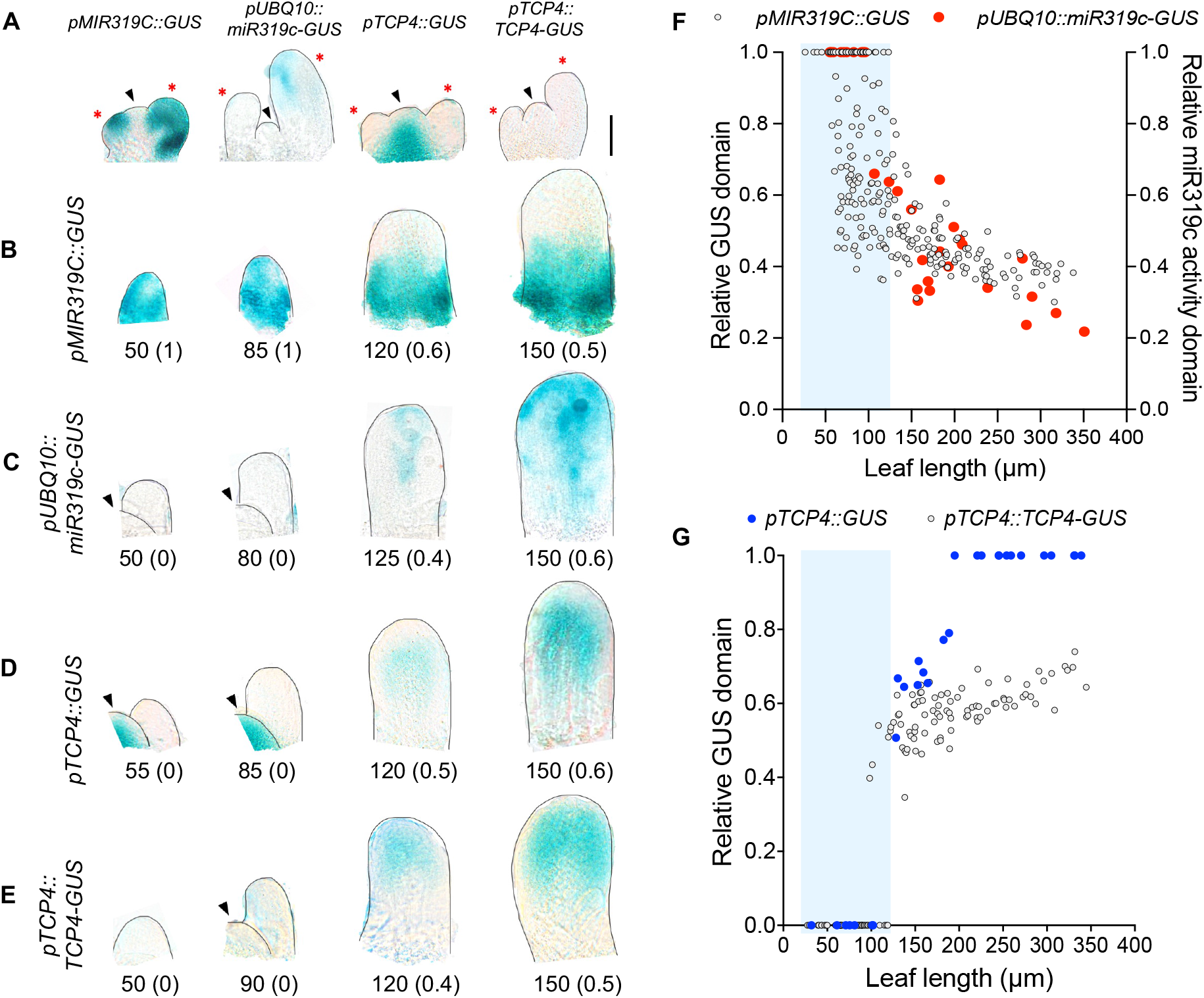
Expression pattern of *MIR319C* and *TCP4* in leaf primordia. **(A)** Bright field images of shoot apices of 10-day-old seedlings of indicated genotypes expressing GUS reporter. Scale bar, 50 μm. Shoot apical meristem (SAM) and leaf primordia are indicated by black arrowheads and red asterisks, respectively. **(B-E)** Bright field images of dissected 7^th^ / 8^th^ leaf primordia expressing indicated *GUS* reporter constructs. Numbers below the images indicate leaf length in μm and numbers within the parentheses indicate the fraction of leaf showing GUS staining along the length axis. SAM is indicated by black arrowheads. **(F)** Scatter plot representation of the relative GUS domain along the length (GUS domain / leaf length) of the 7^th^ / 8^th^ leaf primordia of *pMIR319C::GUS* seedlings (N=242), and the relative miR319c activity domain [(leaf length – GUS domain) / leaf length] of *pUBQ10::miR319c-GUS* seedlings (N=31). **(G)** Scatter plot representation of relative GUS domain along the length of 7^th^ / 8^th^ leaf primordia of *pTCP4::GUS* (N=29), and *pTCP4::TCP4-GUS* (N = 139). The shaded blue boxes mark the leaf primordia with 100% GUS domain in the *pMIR319C::GUS* line **(F)**, or no GUS activity in the *pUBQ10::miR319c-GUS* **(F)**, pTCP4::GUS **(G)** and *pTCP4::TCP4-GUS* **(G)** lines.

To examine whether the pattern of *pMIR319C::GUS* signal reflected miR319c activity, we generated a *pUBQ10::miR319c-GUS* transgenic line where a miR319c-susceptible *GUS* transcript was expressed under the *UBIQUITIN10* promoter (Norris et al., 1993; Geldner et al., 2009). Because *pUBQ10* is ubiquitously and constitutively active in all plant tissues (Li et al., 2019; Zhang et al., 2020), and because the *miR319-GUS* transcript is susceptible to miR319c-mediated degradation, the absence of GUS signal in the *pUBQ10::miR319c-GUS* leaves acted as a readout of mature miR319c activity (**Fig. S3**). No GUS signal was detected till leaves grew to 100 µm (N=10, **Fig. 1C, 1F**), suggesting miR319c activity throughout the primordia. Beyond this stage, GUS signal appeared in distal half of the leaf, indicating a lack of miR319c activity (**Fig. 1C, 1F**). Thereafter, the proportion of GUS domain progressively extended proximally, occupying ∼80% of leaf length when the primordia grew to ∼350 µm long (**Fig. 1F**) and spread nearly throughout the leaf at 1 mm length (**Fig. S2A**).

The above expression analysis suggests an abrupt decrease in *MIR319C* promoter activity in the distal domain of young leaf primordia, with a concomitant loss of mature miR319c activity. Since miR319 degrades *JAW-TCP* transcripts (Palatnik et al., 2003; Nag et al., 2009), its activity is expected to have a negative correlation with JAW-TCP localization. To test this in early leaf primordia, we studied the promoter activity and protein localization of *TCP4*, a *JAW-TCP* member, by analyzing the GUS signal in the *pTCP4::GUS* and the *pTCP4::TCP4-GUS* leaves, respectively (Challa et al., 2019). No *TCP4* promoter activity was detected up to ∼100 µm long leaves (**Fig. 1D, 1G**; in 7 out of 29 primordia studied), a stage when *pMIR319C::GUS* was detected throughout the length in several leaves (**Fig. 1F**). Beyond this stage, *TCP4* promoter activity appeared in the distal half, which spread throughout the leaf by the time the primordia length reached ∼180 µm and beyond (**Fig. 1G, Fig. S2A**). The TCP4-GUS protein signal also remained undetectable till the primordia grew to ∼100 µm long (**Fig. 1E, 1G**; in 52 out of 139 primordia studied), and then appeared in the distal half of the leaf. However, by contrast to the promoter activity, the TCP4-GUS signal domain spread towards the basal region more slowly and reached only ∼70% of the length from the tip when the primordia were ∼350 µm long (**Fig. 1G**). The base remained devoid of *pTCP4::TCP4-GUS* signal even when the leaves grew to 1 mm in length (**Fig. S2A**).

### *MIR319C* promoter activity domain is distally extended in the *JAW-TCP* loss-of-function leaves

The results discussed above demonstrate that *MIR319C* promoter activity is abruptly downregulated in the distal domain of ∼100 µm long leaf primordia with concomitant appearance of TCP4 protein in the same domain (**Fig. 1F, 1G**). To test whether the JAW-TCP proteins are causal to the repression of *MIR319C* promoter, we determined the *pMIR319C::GUS* activity in leaves with reduced *JAW-TCP* level. In wild-type (Col-0;*GR^#1^* and Col-0;*GR^#2^*; **see *Materials* for details**), the average length of the *pMIR319C::GUS* domain was restricted to the basal half (∼45%) of 100-200 µm long leaf primordia (**Fig. 2A, 2B**). However, in the *JAW-TCP* loss-of-function lines *jaw-D* (*jaw-D;GR^#2^*), *pBLS>>miR319a* (*pBLS>>miR319a;GR^#1^*), *tcp2;4;10* (*tcp2;4;10;GR^#1^*) and *tcp2;3;4;10* (Palatnik et al., 2003; Efroni et al., 2008; Challa et al., 2016, Challa et al., 2019) the *pMIR319C::GUS* domain was extended towards the distal region, occupying >65% of the leaf length (**Fig. 2A, 2B**). Out of 152 *JAW-TCP* loss-of-function leaf primordia studied, up to 60% of them showed *pMIR319C::GUS* activity throughout the length at early stages (<120 µm long), whereas <10% of wildtype population showed such widespread GUS expression at comparable growth stage (**Fig. 2C**). The increase in *pMIR319C::GUS* domain persisted at later growth stages when leaves grew to 250-750 µm in length (**Fig. S4A, S4B).** The level of *GUS* transcript also increased up to 2.5-fold in these mutant seedlings compared to the wildtype control (**Fig. S4C**). Further, extended GUS expression was accompanied by 2 to 4-fold increase in pre-*MIR319C* transcript level in the *JAW-TCP* loss-of-function lines (**Fig. 2D**). These results suggest that JAW-TCP activity is required for restricting *MIR319C* promoter activity to the proximal region.

**Figure 2.**
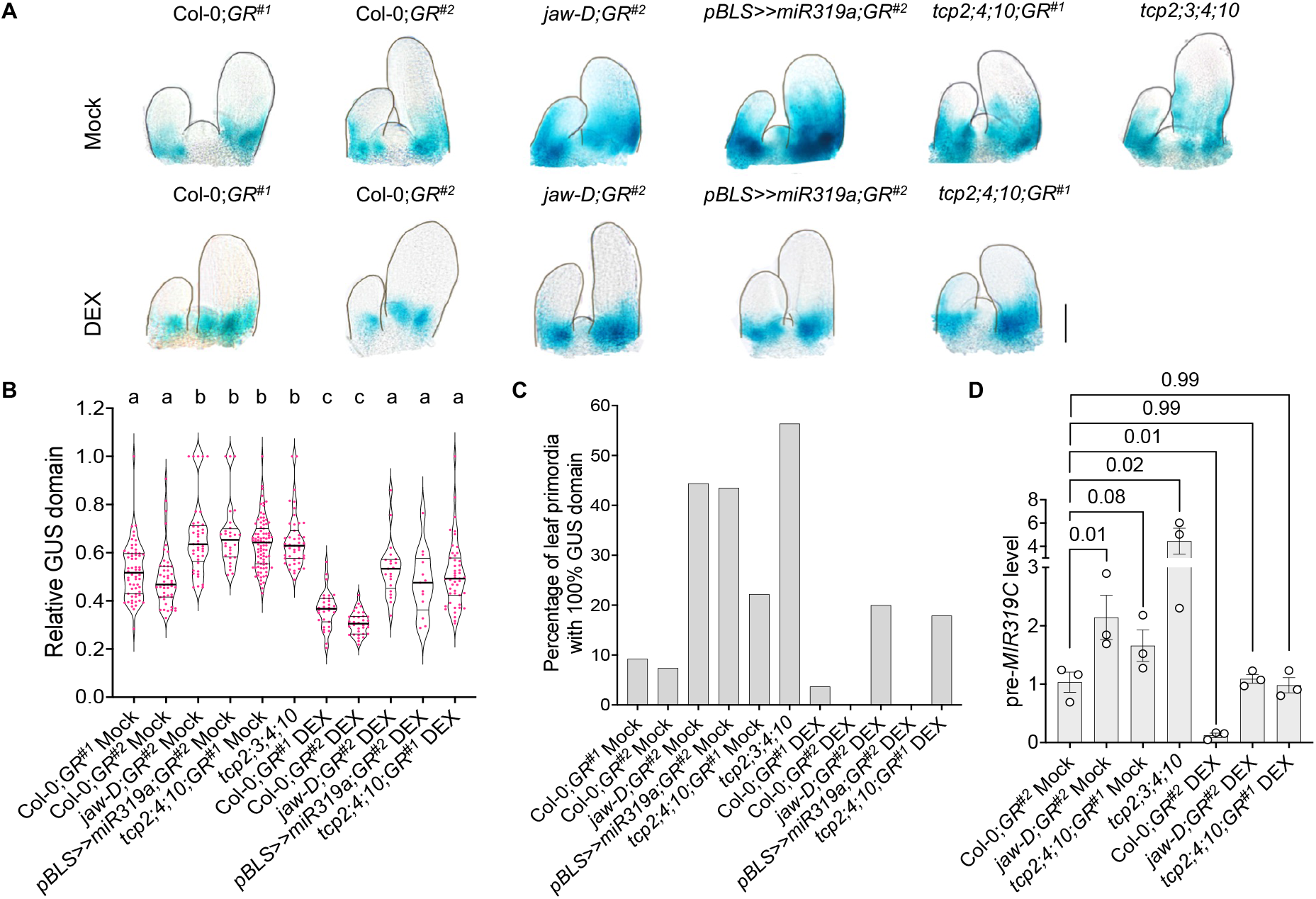
*MIR319C* expression pattern in leaves with altered *JAW-TCPs*. **(A)** Bright field images of 10-day old shoot apices of indicated genotypes processed for GUS activity. Seedlings were grown in the absence (Mock) or constitutive presence (DEX) of 6 µM dexamethasone. Scale bar, 50 μm. **(B)** Violin plot representations of the relative *pMIR319C::GUS* domain in the 100-200 µm long leaf primordia (N=13-87) on 7^th^-8^th^ nodes of seedlings of indicated genotypes treated without (Mock) or with (DEX) 6 µM dexamethasone. Significant differences among samples are indicated by lower-case alphabets. p<0.01, two-way ANOVA, Tukey’s post hoc test was performed. **(C)** Diagram representation of the number of incipient leaf primordia (<120 μm) showing GUS domain throughout the length in genotypes and treatments as indicated along the X-axis. **(D)** Level of the pre-*MIR319C* transcript in 10-day-old seedlings of indicated genotypes treated without (Mock) or with (DEX) 6 µM dexamethasone. Data points from three biological replicates are shown. Error bars indicate SEM. Differences in values among samples are indicated by *p*-values on top of the data points. One-way ANOVA, Dunnett’s post hoc test was performed.

To examine whether JAW-TCP members are sufficient for *pMIR319C* repression, we induced an miR319-resistant form of TCP4 protein (rTCP4-GR; Challa et al., 2016, 2019) which can be activated by administering the glucocorticoid analogue dexamethasone, in the wildtype and in *JAW-TCP* loss-of-function plants. In the wildtype backgrounds Col-0;*GR^#1^* and Col-0;*GR^#2^*, TCP4 induction reduced the *pMIR319C::GUS* domain by restricting it more proximally compared to the uninduced control (**Fig. 2A, 2B, S4A**), accompanied by a ∼2.5-fold reduction in *GUS* transcript level in Col-0;*GR^#2^* (**Fig. S4C**). TCP4 induction also resulted in nearly 6-fold reduction in the pre-*MIR319C* transcript level in Col-0;*GR^#2^* seedlings (**Fig. 2D**). In the three *JAW-TCP* loss-of-function lines where the *pMIR319C* activity domain was distally extended and pre-*MIR319C* transcript level was increased under uninduced condition, TCP4 induction restored the proximal *pMIR319C::GUS* domain (**Fig. 2A, 2B, S4A**), reduced the *GUS* transcript level below the wildtype level (**Fig. S4C**), and decreased the pre-*MIR319C* level to the values similar to that of wildtype (**Fig. 2D**). The *pMIR319C::GUS* domain retracted more proximally also when an inducible, dominant form of *TCP3*, another member of *JAW-TCP* family, was induced in the Col-0;*pTCP3::rTCP3-GR* leaves (**Fig. S4D-F**).

Though *MIR319C* promoter activity was reduced in dexamethasone-treated Col-0;*GR^#1^*, Col-0;*GR^#2^*and Col-0;*pTCP3::rTCP3-GR* leaf primordia (**Fig. 2A, 2B, S4D-F**), it was not completely abolished, perhaps due to the lack of promoter activity of *TCP4*/ *TCP3* at the base of young leaves (**Fig. 1D, 1G**). To test whether the JAW-TCP proteins are capable of repressing *MIR319C* promoter at the extreme leaf base, we generated Col-0;*pMIR319C::rTCP4-GR* and Col-0;*pRPS5A::rTCP4-GR* transgenic lines where rTCP4-GR protein can be induced either in the *MIR319C* expression domain (Nag et al., 2009) or throughout the plant (Weijers et al., 2001), respectively, and analysed *pMIR319C::GUS* expression domain in these leaves. Though under uninduced condition the *pMIR319C::rTCP4-GR*;*pMIR319C::GUS* and the *pRPS5A::rTCP4-GR*;*pMIR319C::GUS* leaves recapitulated the endogenous *MIR319C* promoter activity, rTCP4 induction almost completely cleared the leaf primordia of GUS expression, with only speckles of signal remaining at the junction of leaf and shoot meristem (**Fig. 3A**). Quantification of the average GUS domain in a population of leaf primordia revealed that the degree of reduction in *pMIR319C::GUS* domain in the dexamethasone-treated Col-0;*pMIR319C::rTCP4-GR^#1.2^* and Col-0;*pRPS5A::rTCP4-GR^#1.1^* leaf primordia was nearly 1.7-fold higher than dexamethasone-treated Col-0;*GR^#1^* and Col-0;*GR^#2^* leaf primordia (**Fig. 3B**). Whereas all 31 dexamethasone-treated Col-0;*GR* primordia of length <100 µm studied showed *pMIR319C::GUS* signal at the base, nearly half of *pMIR319C::rTCP4-GR* primordia (19 out of 43) and approximately one-third of *pRPS5A::rTCP4-GR* primordia (8 out of 28) were totally devoid of GUS signal at a similar stage (**Fig. 3C**).

**Figure 3.**
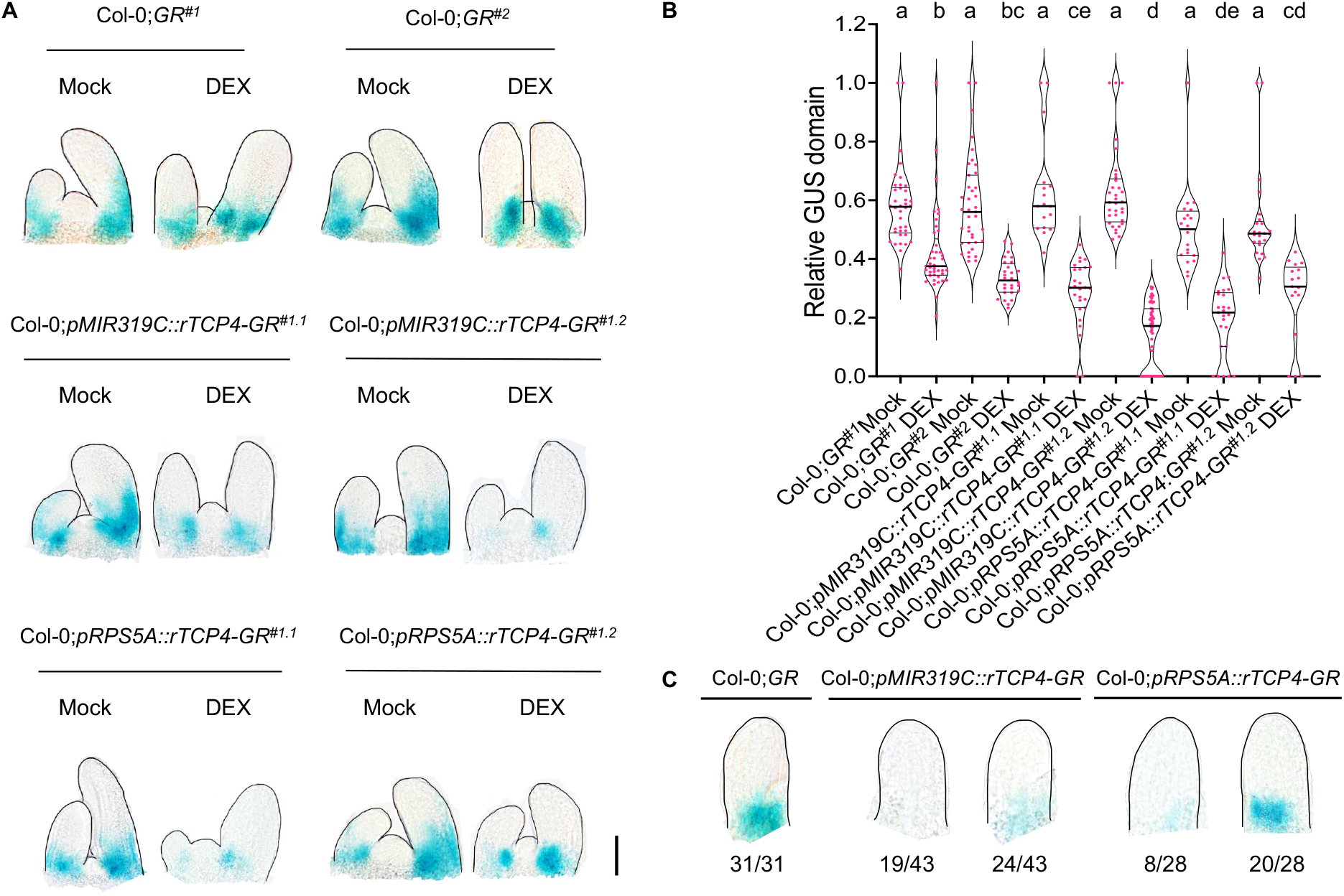
Altered *MIR319C* promoter activity upon ectopic *TCP4* expression in early leaf primordia. **(A)** Bright field images of 10-day old shoot apices of the indicated genotypes expressing *pMIR319C::GUS* reporter constitutively treated with ethanol (Mock) or 6 µM dexamethasone (DEX). Scale bar, 50 μm. **(B)** Violin plots displaying the relative *pMIR319C::GUS* domain in the 50-150 µm long leaf primordia at the 7^th^ and 8^th^ nodes of seedlings of indicated genotypes treated without (Mock) or with (DEX) 6 µM dexamethasone. N=17-41. Significant differences among samples are indicated by lower-case alphabets. p<0.01, two-way ANOVA, Tukey’s post hoc test was performed. **(C)** Bright field images of ∼75 µm long leaf primordia at the 7^th^ or the 8^th^ nodes of seedlings of indicated genotypes (two independent lines combined) expressing *pMIR319C::GUS* reporter constitutively treated with 6 µM dexamethasone. Numbers below the images indicate the number of leaves showing the GUS pattern out of the total leaves analyzed.

In the loss-of-function mutants of *JAW-TCPs*, cells at the leaf base proliferate for longer duration whereas gain-of-function of these genes results in early cell maturity (Efroni et al., 2008; Sarvepalli and Nath, 2011; Challa et al., 2019). Therefore, it can be argued that the perturbation in the *MIR319C* promoter activity described above is a result of altered cell proliferation in the leaf primordia rather than an effect of JAW-TCP activity on *MIR319C* transcription. To discriminate between these two possibilities, we quantified the *pMIR319C::GUS* expression domain in leaves where cell proliferation status is perturbed due to mutations in genes other than *JAW-TCP*s. Though cell proliferation is decreased in the leaves of the *klu-4* mutant and enhanced in the *p35S::ARGOS* plants (Hu et al., 2003, Anastasiou et al., 2007), the *pMIR319C::GUS* domain remained unaltered in 9-day old shoot apices and in leaf primordia compared to wildtype (**Fig. S5A, S5B**), suggesting that the altered *MIR319C* promoter activity described in **Fig. 2** is not merely dependent on the proliferation status of the cells in leaf primordia but is rather an outcome of JAW-TCP activity. This conclusion is further supported by the observation that the average length of early leaf primordia remained unaltered irrespective of the induction of JAW-TCP proteins in the Col-0;*GR^#2^*plants (**Fig. S5C, S5D**).

Together, these results suggest that the JAW-TCP proteins are not only required to restrict the *pMIR319C* activity at the base of Col-0 leaves, but also are sufficient to repress it throughout the leaf primordia when expressed ectopically.

### JAW-TCP proteins are recruited to the *MIR319C* locus

The above results suggest that JAW-TCPs repress *MIR319C* transcription (**Figs. 2, 3**), either directly or by an indirect means. Previous chromatin immunoprecipitation (ChIP)-deep sequencing results suggested that TCP4 protein is enriched at the *MIR319C* locus (Dong et al., 2019) and that pre-*MIR319C* level is 1.7-fold downregulated within 4 hours of TCP4 induction in a microarray experiment (Challa et al., 2016; **Fig. S2B**). Since JAW-TCP proteins are DNA-binding transcription factors, we tested whether they modulate *MIR319C* transcription by directly binding to the *MIR319C* locus. Analysis of *MIR319C* genomic DNA identified the presence of four class II core TCP-binding sequence elements TGGNCC (Schommer et al., 2008; Aggarwal et al., 2010), named here as binding sites 1-4 (BS1-4, **Fig. 4A**). Recombinant 6XHis-TCP4 protein retarded the mobility of radiolabeled oligonucleotide probes containing either of these four sequence elements in an electrophoretic mobility shift assay experiment (**Fig. 4B**). Whereas addition of unlabeled oligonucleotides to the reaction mixture reduced the radioactive signal of the retarded band in a dose-dependent manner, oligos carrying mutations in the consensus sequence (mCP) did not reduce the signal (**Fig. 4B**). These results suggest that TCP4 protein is capable of binding specifically to the *MIR319C* promoter elements in vitro. We independently tested the ability of the JAW-TCPs to bind to these promoter elements in a yeast one-hybrid assay (**Fig. S6**). When a distal upstream regulatory region (URR) containing −2736 to −1866 bp fragment (*p*^distal^) of *MIR319C* locus was used as the promoter to drive the expression of the *HISTIDINE3* (*HIS3*) selection gene in yeast, all five JAW-TCP proteins supported growth of yeast cells in the absence of histidine (**Fig. S6A, S6B**), suggesting that these proteins bind to the TCP4-binding elements present in this URR. By contrast, none of the five JAW-TCP proteins supported yeast growth when *HIS3* was driven by *MIR319C* truncated URR elements from −1900 to −1100 bp (*p*^middle^) or from −1100 to +1 bp (*p*^proximal^) (**Fig. S6A, S6C, S6D**), even though the latter contain three putative TCP-binding elements (BS2-4). Together, these results suggest that JAW-TCPs bind to the distal *cis*-element (BS1) but not to the more proximal elements in the *MIR319C* URR.

**Figure 4.**
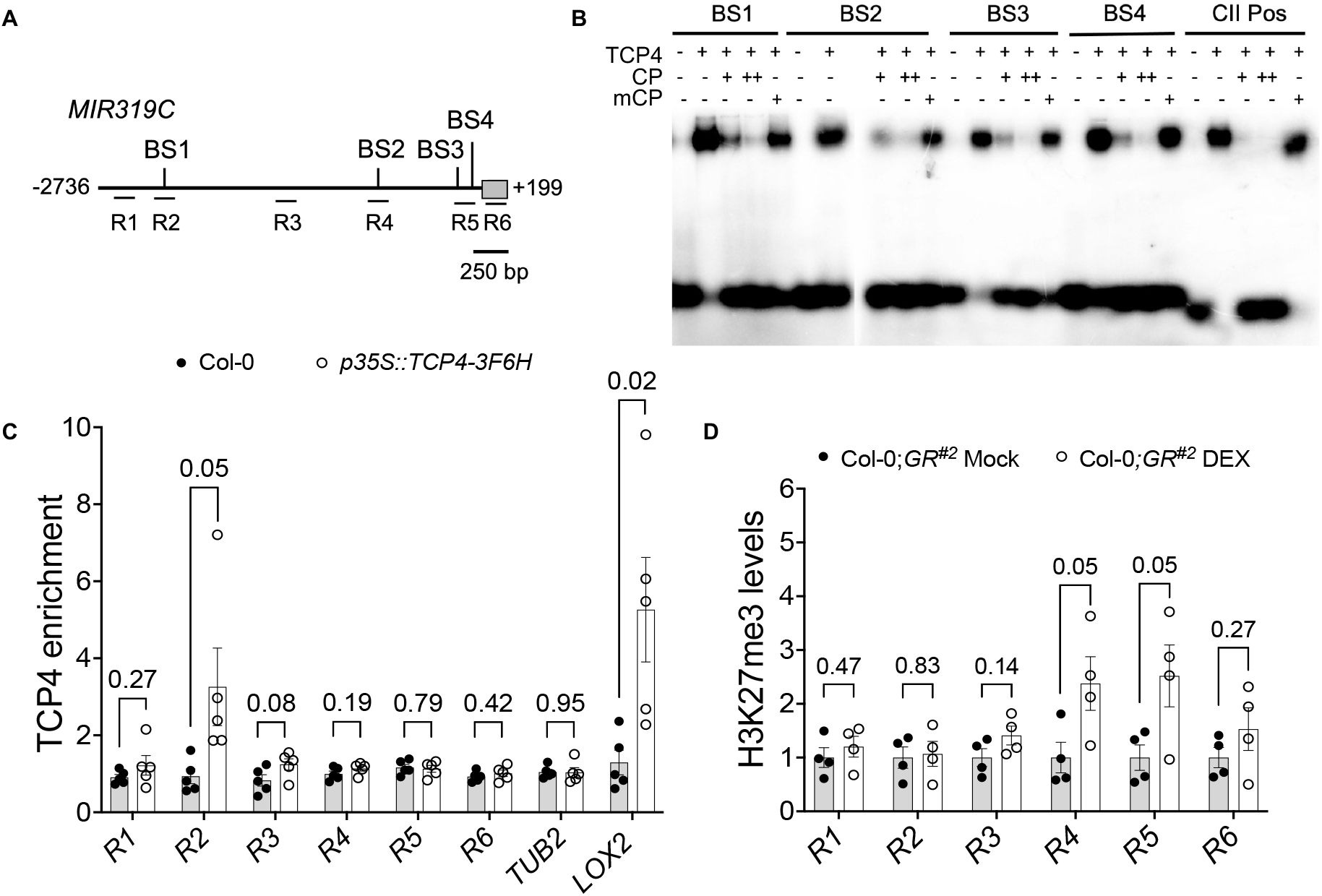
Recruitment of the TCP4 protein on the *MIR319C* locus. **(A)** Schematic representation of the *MIR319C* locus highlighting the consensus TCP4-binding sites (BS1-BS4) in the upstream regulatory region. R1-R6 correspond to the regions amplified in the ChIP-qPCR experiment shown in **(C)** and **(D)**. Grey box corresponds to the 199 bp-long pre-*MIR319C* transcript. **(B)** Image of the autoradiogram of an EMSA experiment showing the retardation of radiolabeled oligonucleotide probes corresponding to BS1-BS4 by recombinant the 6XHis-TCP4 protein (TCP4). A labeled probe containing class II consensus sequence GTGGNCCC (CII Pos) served as the positive control (Aggarwal et al., 2010). The + and the – symbols indicate the presence and the absence of the indicated reagents, respectively. Un-labelled probes with wild-type (CP) or mutated (mCP) sequence were added in 10-fold (+) or 25-fold (++) excess. **(C)** Fold enrichment of the fragments corresponding to R1-R6 regions shown in (**A**) precipitated by anti-FLAG antibody in the ChIP experiment with chromatins isolated from 9-day old Col-0 or *p35S::TCP4-3F6H* seedlings (Kubota et al., 2017). Fold enrichment of TCP4-FLAG was calculated relative to IgG. *β-TUBULIN* (*TUB2*) and *LIPOXYGENASE2* (*LOX2*) (Challa and Rath et al., 2021) promoter regions were used as negative and positive controls, respectively. Error bars represent SEM of five biological replicates, each performed in technical duplicates. Pairwise comparisons were performed using unpaired *t*-test. *p*-values are shown above the data points. **(D)** Fold enrichment of the fragments corresponding to R1-R6 regions precipitated by anti-histone H3K27me3 antibody in the ChIP experiment with chromatins isolated from 9-day old Col-0;*GR^#2^* seedlings grown in the absence (Mock) or presence (DEX) of 6 µM dexamethasone. Fold change in histone H3K27me3 levels is represented as %age input and normalized to values from respective mock-treated Col-0;*GR^#2^* controls. Error bars represent SEM of four biological replicates, each performed in technical duplicates. Pairwise comparisons were performed using unpaired *t*-test. *p*-values are shown above the data bars.

To determine whether JAW-TCPs bind to the *MIR319C* URR elements in planta, we carried out a ChIP experiment. Quantitative PCR (qPCR) analysis of the chromatin fragments prepared from the *p35S::TCP4-3F6H* seedlings (Kubota et al., 2017) and precipitated with anti-FLAG antibody using primers against regions R1-R6 (**Fig. 4A**) showed TCP4 enrichment at the R2 region that encompasses the BS1 sequence element (**Fig. 4C**). However, no such enrichment was detected in the URR regions corresponding to R2-R6 sites, suggesting that TCP4 is recruited specifically to the BS1 region of the *MIR319C* promoter.

All JAW-TCP members are known to bind directly to the *cis*-elements of their target genes and activate their transcription (Aggarwal et al., 2010; Challa et al., 2016, 2019; Vadde et al., 2018, 2019). However, their transcription repression function, as suggested above for the *MIR319C* locus (**Figs. 2, 3**), is not explicitly reported. Therefore, the molecular mechanism of *MIR319C* suppression by JAW-TCPs remains speculative. JAW-TCPs are known to form complexes with transcription factors which recruit histone methyltransferases and repress their target gene transcription (Li et al., 2012; Lodha et al., 2013). It is, hence, possible that the JAW-TCPs recruit histone-modifying enzymes to suppress *MIR319C* transcription. In addition, CURLY LEAF (CLF) and SWINGER (SWN), two of the three histone methyltransferases of the polycomb repressive complex2 (PRC2) responsible for tri-methylation of histone H3 at the lysine 27 position (Shen et al., 2021), were significantly enriched on the *MIR319C* locus in a previously reported ChIP-seq study (Shu et al., 2019). Hence, we examined the relative level of histone H3K27me3 modification, a repression mark, on *MIR319C* locus. Upon dexamethasone induction in 9-day old Col-0;*GR^#2^* seedlings, histone H3K27me3 level was 2.5-fold higher at the R4 (−500 bp) and the R5 (−150 bp) regions of *MIR319C* compared to the uninduced control (**Fig. 4D**), suggesting that TCP4 is capable of enhancing the repressive state of *MIR319C* chromatin. The histone H3K27me3 mark on the chromatin of the *JAW-TCP* loss-of-function line *tcp2;3;4;10*, however, was not reduced compared to the Col-0 value (**Fig. S6E**), perhaps due to the presence of other four class II TCP members (*TCP5*, *13*, *17* & *24*) in this mutant. The level of histone H3K14Ac activation mark also remained similar in wildtype and *tcp2;3;4;10* chromatin at the *MIR319C* locus (**Fig. S6F**).

### Compromised JAW-TCP binding leads to increased *MIR319C* promoter activity

If repression of *MIR319C* transcription (**Figs. 2, 3**) is mediated by JAW-TCP binding to its promoter elements (**Fig. 4**), then mutating the TCP4-binding *cis*-elements in the *MIR319C* promoter region is expected to increase its activity. To test this, we generated a *mutpMIR319C::GUS* transgenic line where the reporter gene was expressed under *MIR319C* promoter carrying point mutations at all four class II TCP-binding sites that abolished TCP4-binding (**Fig. 4B, 5A**). When this line was crossed to Col-0;*GR^#1^*, the resulting *mutpMIR319C::GUS* x Col-0;*GR^#1^* F1 leaves expressed GUS signal in the proximal 70% of length, similar to what was observed in the *pMIR319C::GUS^#2^*x Col-0;*GR^#1^* negative control (**Fig. 5B, 5C, S7A**). Further, activation of the dominant rTCP4-GR in the *mutpMIR319C::GUS* x Col-0;*GR^#1^* F1 leaves reduced the GUS domain to the same extent as was observed in the *pMIR319C::GUS^#2^*x Col-0;*GR^#1^* leaves where the TCP-binding sites were not mutated (**Fig. 5B, 5C, S7A**). These results suggest that mutation in the TCP-binding elements fails to enhance *MIR319C* transcription, at least in the Col-0 background. To examine whether this failure is due to the presence of endogenous level of JAW-TCP proteins, we reduced the *JAW-TCP* level in the *mutpMIR319C::GUS* plants by crossing it to the *jaw-D;GR^#1^* line. The resulting *mutpMIR319C::GUS* x *jaw-D;GR^#1^* leaves also showed GUS activity similar to the *pMIR319C::GUS* x *jaw-D;GR^#1^* control leaves, possibly because of a single copy of the *jaw-D* allele. However, induction of rTCP4-GR in the *mutpMIR319C::GUS* x *jaw-D;GR^#1^*leaves did not restrict the GUS domain more proximally compared to the uninduced control, though GUS restriction was observed when *MIR319C* promoter was not mutated (**Fig. 5B, 5C, S7A**). This suggests that, at reduced level of *JAW-TCP* transcripts, TCP4 fails to repress *MIR319C* transcription if its binding to the promoter elements is compromised.

**Figure 5.**
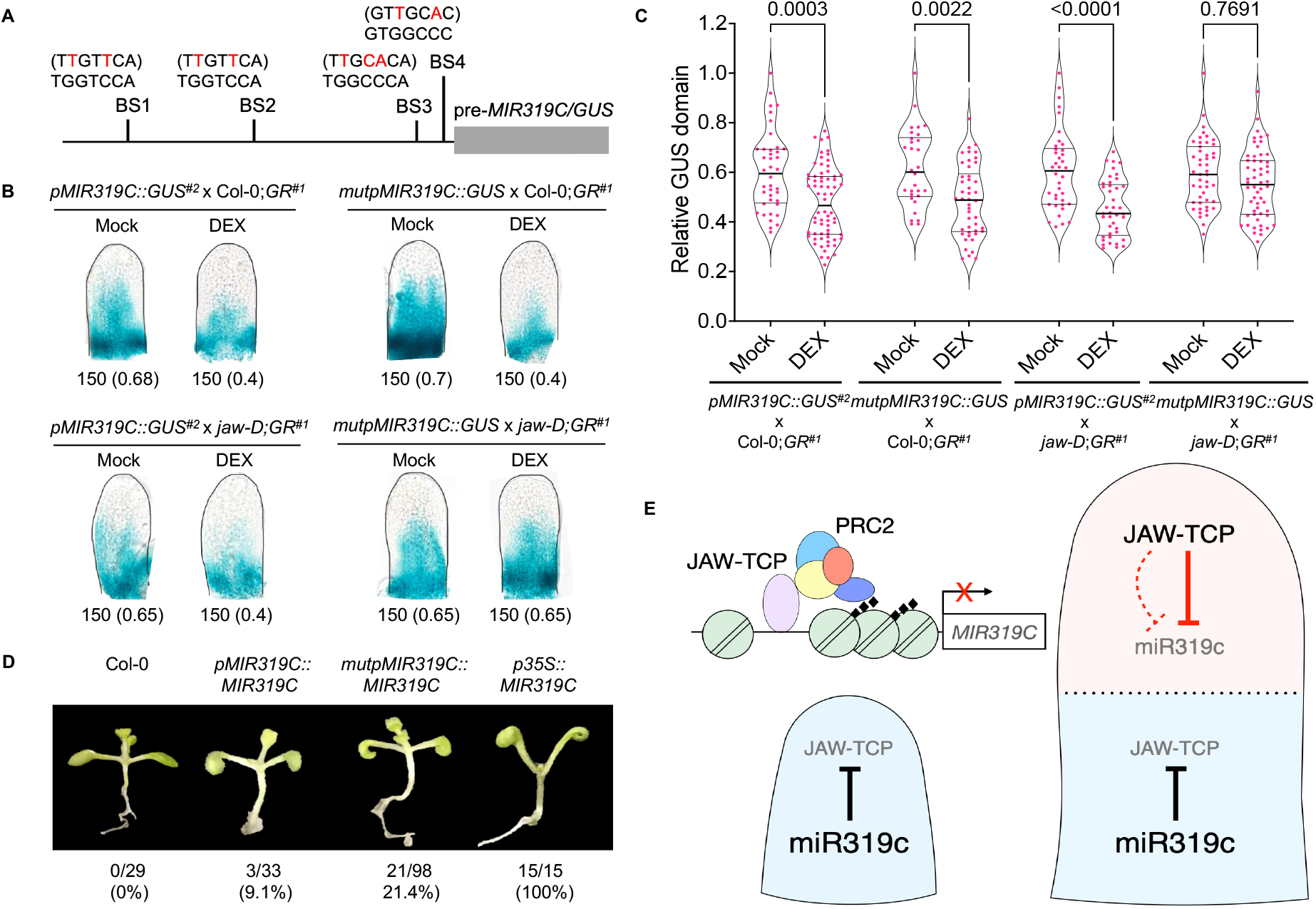
Altered JAW-TCP binding to the *MIR319C* upstream regulatory region with mutated TCP-binding sites. **(A)** Schematic representation of the *pMIR319C::MIR319C/GUS* construct with mutated bases on the core TCP-binding sites indicated in red. The pre-*MIR319C*/*GUS* region is indicated by grey box. **(B)** Bright field images the 7^th^ leaf primordia of 10-day old seedlings grown in the absence (Mock) or presence (DEX) of 6 µM dexamethasone and processed for GUS activity. Numbers below the images indicate leaf length in µm and relative GUS domain (in parentheses). **(C)** Violin plot representations displaying the relative *GUS* domain in 50-350 µm long 5^th^-8^th^ leaf primordia (N=30-68) from 10-day old seedlings grown in the absence (Mock) or presence of 6 µM dexamethasone (DEX). Significant differences among samples are indicated by *p*-values on top of the plots. Two-way ANOVA, Tukey’s post hoc test was performed. **(D)** Representative images of 9-day old Col-0 and transgenic (T1) seedlings expressing *pMIR319C::MIR319C*, *mutpMIR319C::MIR319C*, or *p35S::MIR319C* constructs. Numbers and percentages of transgenic lines showing epinastic cotyledons out of the total number of independent insertion lines analyzed here are shown below the images. Col-0 and *p35S::MIR319C* lines were used as negative and positive controls, respectively. **(E)** Schematic representation of a proposed double-negative feedback loop between *MIR319C* and *JAW-TCPs*. The *MIR319C* promoter is active (light blue) throughout the incipient leaf primordium (left panel) prior to the activation of *JAW-TCP* transcription, where the mature miR319c ensures that no basal-level JAW-TCP proteins are accumulated by degrading their transcripts (solid black **T** line). As the primordium grows out of the apical meristem, *JAW-TCP* transcription is activated towards the distal region (light red on the right panel) by activators that are yet unknown. The level of JAW-TCP proteins that accumulate at the distal region despite the activity of miR319c, bind to the upstream regulatory region of *MIR319C* and trigger the silencing of its transcription (solid inverted red **T** line), possibly by recruiting the PRC2 chromatin silencing components, thereby ensuring the build-up of JAW-TCP proteins. JAW-TCPs may also suppress *MIR319C* transcription by indirect mechanisms (dotted inverted red **T** line), such as activation of a repressor protein or inhibition of an activator protein by physical interaction. Mature miR319c activity continues to suppress the basal level JAW-TCP accumulation at the proximal region. Thus, a double-negative feedback interaction between miR319 and JAW-TCPs divides the uniformly proliferating incipient leaf primordia into two distinct growth domains – the distal zone of cell differentiation and the proximal zone of cell proliferation, separated by a diffused boundary (broken horizontal line). Black diamonds represent H3K27me3 marks, green circles represent nucleosomes, arrow indicates transcriptional activation.

To test the phenotypic consequences of the mutations in the TCP4-binding sites on the *MIR319C* URR, we generated a *mutpMIR319C::MIR319C* transgenic line where pre-*MIR319C* was expressed under the mutated *MIR319C* promoter (**Fig. 5A**). It has been demonstrated that plants overexpressing miR319 show cotyledon epinasty and larger rosette leaves with serrated and wavy margin (**Fig. 5D, S7B**; Palatnik et al., 2003; Challa et al., 2016). Nearly 21% of 98 *mutpMIR319C::MIR319C* independent transgenic seedlings exhibited cotyledon epinasty in the T1 generation, as opposed to 9% of 33 *pMIR319C::MIR319C* independent lines where pre-*MIR319C* was expressed under endogenous promoter (**Fig. 5D**), suggesting that binding of JAW-TCP proteins to the *pMIR319C cis*-elements in Col-0 reduces its phenotypic effects. However, the *mutpMIR319C::MIR319C* rosette leaves grew to the same average size at maturity as the *pMIR319C::MIR319C* leaves (**Fig. S7B, C**), indicating that the mutations introduced in the *MIR319C URR* affected embryonic leaves only and not rosette leaves. Together, the above results indicate that JAW-TCPs proximally restrict *MIR319C* expression domain at least partly by binding to the *cis*-elements in its upstream regulatory region.

## Discussion

Leaf morphogenesis in Arabidopsis is characterized by a brief period of cell proliferation in early primordia, followed by differentiation and expansion (Donnelly et al., 1999; Andriankaja et al., 2012). Differentiation is initiated at the distal end of a young primordium, while cell proliferation continues in the proximal domain. How growth in a uniformly dividing incipient primordium is patterned along the proximal-distal axis remains unclear. A double-negative feedback loop (DNFBL) between a growth activator and a repressor would be a minimalistic proposal to establish this growth polarity (Tsang et al., 2007; Cai et al., 2013). The miR319c-JAW-TCP module plays a key role in leaf morphogenesis wherein the miR319c does not allow the accumulation of *JAW-TCP* transcripts at the base (Palatnik et al., 2003; Nag et al., 2009; Alvarez et al., 2016; Challa et al., 2019). However, how *MIR319C* expression is excluded from the distal domain leading to the initiation of leaf maturation is not known. Here we provide evidence that the JAW-TCP proteins repress *MIR319C* transcription in the distal zone of a leaf primordium, thus forming a DNFBL between them (**Fig. 5E**). According to this model, miR319c is active throughout the incipient leaf primordia (**Fig. 1C, 1F**), perhaps because *MIR319C* transcription is activated by a yet-unknown leaf-specific factor during organ initiation (**Fig. 1A**). Although the *JAW-TCP* promoter activity is not detected at this stage (**Fig. 1D**), ubiquitous miR319c activity would eliminate any basal level of JAW-TCP transcripts and hence their protein products (**Fig. 1E, 1G**), thus ensuring uniform growth and proliferation throughout the primordia. Transcription of *TCP4* and its homologs are activated, perhaps by unknown distal-specific factors, in primordia of 100-150 µm length (**Fig. 1D, 1G**), leading to repression of *MIR319C* promoter activity. The miR319c activity, however, continues at the basal region (**Fig. 1B, 1C, 1F**), thus dividing the primordium into two domains of contrasting proliferating status (**Fig. 5E**). Activation of *JAW-TCPs* and concomitant abrupt decrease in miR319c activity (**Fig. 1F and 1G**) resembles a typical miRNA-TF DNFBL, wherein a trigger factor increases TF activity and divides a homogenous field of miRNA expressing cells into two distinct domains where miRNA and TF repress each other (Li and Carthew, 2005; Cai et al., 2013).

Multiple mutants with reduced JAW-TCP activity exhibited distally expanded domain of *MIR319C* expression and had increased pre-*MIR319C* levels (**Fig. 2A-D**). On the other hand, *MIR319C* expression domain was restricted more proximally in mutants with increased TCP3/ TCP4 level (**Fig. S4D-F**). In addition, ectopic TCP4 activity throughout the initiating leaf primordia was sufficient to completely abolish *MIR319C* expression (**Fig. 3**). This suggests that JAW-TCPs redundantly restrict *MIR319C* expression domain towards the base of the leaf primordia. The recession of *MIR319C* expression in the distal region of the primordia was, however, only delayed but not abolished in mutants with reduced JAW-TCP activity (**Fig. 2A, S4A, S4B**), supporting their role as heterochronic factors (Efroni et al., 2008) and suggesting that *MIR319C* expression is perhaps repressed by additional factors. It would be interesting to investigate if the differentiation factors belonging to the *NGATHA* and *KNOX-II* gene families, whose expression and functions overlap with *JAW-TCPs* (Alvarez et al., 2016; Challa and Rath et al., 2021), downregulate *MIR319C* expression in distal leaf primordia. The altered domain of *MIR319C* expression seen in *JAW-TCP* mutants is not due to the changes in the proliferation status of the cells. Multiple studies report that proliferation occurs throughout the leaf primordia up to the stage of 250 μm length (Kazama et al., 2010; Ichihashi et al., 2011; Andriankaja et al., 2012). The *MIR319C* expression gets restricted to the base much earlier, when the primordium length is 150 μm or less, suggesting that downregulation of *MIR319C* in the distal region occurs before the onset of the switch from proliferation to differentiation. Additionally, *MIR319C* expression remains unperturbed in the leaves of *klu-4* and *p35S::ARGOS* mutants where the duration of cell proliferation is altered independent of *JAW-TCPs* (**Fig. S5A and S5B**).

We have provided evidence that JAW-TCPs bind to the URR elements of *MIR319C*, downregulate its expression, and induce the repression marks on its chromatin. It is possible that all these processes are linked, and that JAW-TCPs repress *MIR319C* transcription by modifying its chromatin structure, possibly by recruiting histone modifying-enzymes such as the PRC2 members (Lodha et al 2013). Though TCP4 was distinctly enriched at the distal URR of *MIR319C* spanning BS1 in planta (**Fig. 4C**), the histone H3K27me3 repressive marks were enhanced at the regions proximal to the site of TCP4 enrichment (**Fig. 4D**). JAW-TCPs are known to form dimers not only among themselves (Aggarwal et al., 2010; Danisman et al., 2013), but also with transcription factors belonging to diverse gene families (Li et al., 2012, Rubio-Somoza et al., 2014). Hence it is possible that JAW-TCPs bind at the distal URR and interacts with TCP homologues and other TFs that may bind to the proximal *cis*-elements. This homo and heterodimer formation is expected to induce looping of the chromatin, resulting in enhanced recruitment of histone methyltransferase complex at the proximal URR and lead to *MIR319C* repression (**Fig. 5E**; Li et al., 2012; Lodha et al., 2013; Ouyang et al., 2020). Examples of such chromatin looping-mediated regulation of gene expression have been reported in several species including Arabidopsis, maize, and rice (Cao et al., 2014; Wang et al., 2019; Ouyang et al., 2020). Currently, no other reports demonstrate chromatin modification as a mechanism for JAW-TCP*-*mediated repression of its target genes, even though circumstantial evidence suggests this possibility. JAW-TCPs redundantly form complexes with AS2 to directly bind and repress the expression of *KNOX-I* genes *BP* and *KNAT2* (Li et al., 2012). Another study shows that AS2 physically interacts and recruit histone methyltransferases of the PRC2 family onto *BP* and *KNAT2* promoters to repress their expression (Lodha et al., 2013). Also, TCP4 interacts with the SWI/SNF chromatin remodeling ATPase BRAHMA (BRM) and together promote leaf maturation by modifying cytokinin responses (Efroni et al., 2013). Hence it is possible that chromatin modification is the primary mechanism of JAW-TCP-mediated regulation of its downstream target genes. Interestingly, TCP4 failed to downregulate the activity of *MIR319C* promoter with mutated *cis*-elements only in the *jaw-D* leaf primordia but not in wild-type, suggesting that TCP4 requires other JAW-TCPs for *MIR319C* repression. However, the lack of JAW-TCP binding did not further extend the *MIR319C* expression domain both in the Col-0 and *jaw-D* backgrounds (**Figs. 5B and 5C**), perhaps indicating the existence of an indirect pathway for JAW-TCP-mediated *MIR319C* repression.

Our results support a model where the miR319c-JAW-TCP DNFBL divides a primordium with homogenous miR319c activity into one where the miR319c and JAW-TCP activities are spatially separated and maintained during subsequent stages of growth (**Fig. 5E**). *MIR319C* expression is seen at the earliest stages of primordia, while the distal specific expression of *TCP4*, and possibly other *JAW-TCPs* leads to stable repression of *MIR319C* via chromatin modification and indirect mechanisms, thus restricting the domain of miR319c activity to the base of the primordia at a later stage (∼150 μm long). JAW-TCP-mediated downregulation of *MIR319C* expression in turn amplifies *JAW-TCP* transcript and establishes domain of high JAW-TCP activity in the distal region. However, *MIR319C* expression is maintained at the base even during the later stages when the *TCP4* promoter is active throughout the leaf primordia, possibly by a persistent activation mechanism. Modelling and experimental evidence suggest that miRNA-TF DNFBLs can promote switching between a high TF and a high miRNA expression state using minimal inputs to regulate cell fate transitions during organ development (Fazi et al., 2005; Johnston et al., 2005; Cai et al., 2013). The DNFBL between miR319 and JAW-TCPs can be established with minimal inputs such as an initial trigger for onset of *JAW*-*TCPs* transcription at the distal end and a strong activator of *MIR319C* expression at the base. Even a relatively weak transcriptional input for *JAW-TCPs* could be translated into a stable *MIR319C* repression due to the positive reinforcement nature of the feedback loop. Detailed kinematic studies during early stages of leaf growth suggests the abrupt nature of the transition of distal cells into state of differentiation (Andriankaja et al., 2012). The stable *MIR319C* repression could quickly ensure the uniformity of JAW-TCP levels in distal cells which in turn would allow them to coherently transition to differentiation. Hence, miR319-JAW-TCP DNFBL could be a part of the gene regulatory network involved in the establishment of cell proliferation patterns in the leaf primordia.

Currently, we do not know how *MIR319C* expression is initiated and later maintained in the proximal region of the leaf primordia. Also, the factors that activate *JAW-TCP* expression in the distal region are not known. Thus, determining the activators of *MIR319C* and *JAW-TCP* transcription, and additional factors for the distal *MIR319C* repression will provide insights into a more complete genetic network that establishes the distinct domains of miR319 and JAW-TCP activity in an otherwise homogenous field of cells with high *MIR319C* expression in early leaf primordia.

## Materials and Methods

### Plant materials

*Arabidopsis thaliana* ecotype Columbia-0 (Col-0) was used as the wild-type control. All the lines used in this study are in Col-0 genetic background. The genotypes used in this study are described in **Table S1**. The genotype *pBLS>>miR319a* (Originally in L*er*; Efroni et al., 2008) was brought into the Col-0 background by back crossing three times. *tcp2;3;4;10* was generated by crossing *tcp2;4;10* (Challa et al., 2016) and *tcp3;4;10* (Koyama et al., 2010) and confirmed by genotyping in the F3 generation. *pMIR319C::GUS^#1^*(Nag et al., 2009) was established in various mutant backgrounds by crossing and was confirmed by GUS assays in the F3/F4 generation.

### Plant growth condition and chemical treatments

Seeds were surface-sterilized with 0.05% SDS containing 70% ethanol solution for 10 min, followed by wash with 100% ethanol twice and sown on 1X MS medium (PT011, HiMedia Laboratories, India) containing 0.025% ethanol (Mock) or 6 μM dexamethasone (DEX) (D4902, Sigma-Aldrich, USA). After sowing, seeds were kept for stratification for three days in the dark at 4°C and then shifted to a plant growth chamber (I-41LL, Percival Scientific, USA) set at 22°C, 100 μmol/m^2^ light intensity and 16 h light/8 h dark light regime. 14-day old seedlings were transplanted onto a soil mixture containing Soilrite and Perlite (Keltech Energies, India) in a 3:1 ratio. Plants were grown till maturity in growth rooms with the conditions as described for growth chamber.

### DNA constructs and transgenic line generation

For the generation of *pUBQ10::miR319c-GUS* construct, a 1.5 kb long fragment corresponding to *UBQ10* upstream regulatory region (URR) ending with the start codon was amplified by PCR using Col-0 genomic DNA as template. A 20-nucleotide sequence, complementary to the mature miR319c, was incorporated into the reverse primer used for the amplification of *pUBQ10*. The *pUBQ10::miR319c* fragment was cloned into modified *P4P1R* entry vector (a kind gift from Ram Yadav, IISER Mohali, India) by gateway BP reaction (11789020, Invitrogen, USA) to generate the *pUBQ10::miR319c-P4P1R* construct. *pUBQ10::miR319c* was subcloned into the *R4L1pGWB532* destination vector (Gift from Tsuyoshi Nakagawa, Shimane University, Japan) by LR reaction (11791020, Invitrogen, USA) to generate the final *pUBQ10::miR319c-GUS* construct.

For the generation of *pTCP3::rTCP3-GR* construct, *TCP3* ORF was first PCR-amplified using Col-0 cDNA, digested with *BamH*I*-Pst*I, and then subcloned into *pGAD424*. To generate miR319-resistant *TCP3*, synonymous point mutations were introduced into *TCP3-pGAD424* by inverse PCR (Palatnik et al., 2003) to generate *rTCP3-pGAD*. The *GR* fragment was cleaved out from *pTCP4::rTCP4-GR* (Challa et al., 2016) by *BamH*I*-Nco*I digestion and cloned in-frame with *rTCP3* at its C-terminus in *pCAMBIA 1390* to generate *rTCP3-GR-pCAMBIA 1390*. A 2.5 kb URR of *TCP3* locus (Koyama et al., 2007) was digested from the *pTCP3-BJ36* construct (Alvarez et al., 2016) using *BamH*I-*Pst*I enzymes and cloned into *pCAMBIA 1300* vector. The *TCP3* promoter was further moved into *pCAMBIA 1390* using *BstX*I-*Pst*I. *rTCP3-GR* fragment from *rTCP3-GR-pCAMBIA 1390* was digested with *Sal*I-*Nco*I and cloned downstream of *pTCP3* in *pTCP3-pCAMBIA 1390* to generate *pTCP3::rTCP3-GR* construct. For the generation of *pTCP4::rTCP4-GR* construct, *rTCP4-GR* fragment from *rTCP4-GR-KS* (Challa et al., 2016) was cloned into *pCAMBIA 1390* to generate *rTCP4-GR-pCAMBIA 1390*. *rTCP4-GR* was digested with *Sal*I-*Nco*I and cloned downstream of 2.16 kb *TCP4* promoter in the *pTCP4-pCAMBIA 1390* (Challa et al., 2016) to generate *pTCP4::rTCP4-GR*.

For the generation of *pMIR319C::MIR319C*, *mutpMIR319C::MIR319C*, *pMIR319C::GUS*, *mutpMIR319C::GUS*, *pMIR319C::rTCP4-GR*, and *pRPS5A::rTCP4-GR* constructs, a 2.7 kb genomic fragment corresponding to the *MIR319C* URR from the start of pre-*MIR319C* was amplified from Col-0 genomic DNA using Phusion High Fidelity DNA polymerase (M0530, New England Biolabs, USA) and cloned into *P4P1R* to generate the *pMIR319C-P4P1R* construct. *rTCP4-GR* fragment was amplified from *pTCP4::rTCP4-GR pCAMBIA 1390* construct (Challa et al., 2016) and introduced into the modified *pDONR221* donor vector (Gift from Ram Yadav, IISER Mohali, India) by gateway BP reaction. *pMIR319C* and *rTCP4-GR* entry clones were confirmed by sequencing and recombined into the destination vector *R4pGWB507* (Gift from Tsuyoshi Nakagawa, Shimane University, Japan) by gateway LR reaction to generate the *pMIR319C::rTCP4-GR* construct. A similar strategy was used to generate the *pRPS5A::rTCP4-GR* construct.

For the generation of *mutpMIR319C* construct, we utilized a PCR-based site directed mutagenesis approach. Briefly, forward, and reverse primers carrying mutations in class II TCP binding site 1 (mBS1) was first used in a PCR reaction with 2.7 kb *pMIR319C-P4P1R* template using Phusion DNA polymerase. Post PCR reaction, the product was digested with *Dpn*I (R0176, New England Biolabs, USA) and transformed into *E. coli* and selected on LB agar containing 100 mg/l ampicillin. Plasmids were purified from resulting colonies using QIAprep Spin Miniprep Kit (Qiagen, Germany). The final *mutpMIR319C-P4P1R* construct was generated by introducing mBS2, mBS3 and mBS4 sequentially into *mBS1pMIR319C-P4P1R*, *mBS12pMIR319C-P4P1R* and *mBS123pMIR319C-P4P1R* templates and confirmed by sequencing. A 199 bp fragment corresponding to pre-*MIR319C* was amplified from Col-0 genomic DNA using Phusion High Fidelity DNA polymerase and cloned into the *pDONR221* donor vector by BP reaction. The entry clones *mutpMIR319C-P4P1R* and pre-*MIR319C*-*pDONR221* were recombined into the *R4pGWB507* destination vector by LR reaction to generate the *mutpMIR319C::MIR319C* construct. *pMIR319C-P4P1R* and pre-*MIR319C*-*pDONR221* entry clones were recombined into the *R4pGWB507* destination vector by LR reaction to generate the *pMIR319C:::MIR319C* construct. Similar approach was undertaken to generate *p35S::MIR319C* and *pRPS5A::MIR319A* constructs. The *pMIR319C::GUS* and *mutpMIR319C::GUS* constructs were generated by recombining *pMIR319C-P4P1R* and *mutpMIR319C-P4P1R* entry clones into *R4L1pGWB532* destination vector, respectively, by LR reaction. Sequence information of primers used to generate the constructs are provided in the **Table S2**.

While the *pMIR319C::rTCP4-GR* and *pRPS5A::rTCP4-GR* constructs were integrated into the *pMIR319C::GUS*;Col-0 background, all other binary constructs were integrated into Col-0 background by the *Agrobacterium tumefaciens*-mediated floral dip method (Clough and Bent, 1998) except Silwet L-77 was used at a concentration of 50 μl/litre. Following transformation, seeds were harvested and screened for the ability to grow on MS media in presence of 20 mg/l hygromycin. At least five independent lines were established for each construct and confirmed by genotyping and phenotyping in the T3 generation. Information on PCR-based genotyping primers are provided in the **Table S2**.

### β-glucuronidase (GUS) assay

GUS assays were performed as described earlier (Challa et al., 2016). Briefly, samples were collected in ice-cold 90% acetone, incubated at room temperature for 20 min, followed by washing with ice-cold staining buffer (50 mM sodium phosphate, pH 7.0, 0.2% Triton X-100, 5 mM potassium ferrocyanide, and 5 mM potassium ferricyanide). For *pUBQ10::miR319c-GUS*, staining buffer containing 10 mM potassium ferrocyanide and 10 mM potassium ferricyanide was used. Fresh staining buffer was added with 2 mM X-Gluc (R0851, Thermo Scientific, USA) and incubated for 4-6 hours (for *pTCP4::GUS; pTCP4::TCP4-GUS, pUBQ10::miR319c-GUS* and *pMIR319C::GUS^#2^*) or overnight (for *pMIR319C::GUS^#1^*) at 37°C in dark. Staining buffer was replaced with 70% ethanol, and 70% ethanol was changed 3-4 times in regular intervals to clear the tissue. Shoot apices and leaf primordia were dissected and mounted on a glass slide containing lactic acid under the microscope and were imaged using Olympus BX51 trinocular DIC microscope fitted with ProgResC3 camera and ProgResCapture Pro2.6 software (Jenoptik, Germany) with the following settings, exposure-65ms, brightness-10 units, contrast-35 units, colour temperature-5000K, Image resolution-(1040 x 770 2X Bin), while keeping all other settings in default mode.

### GUS domain and leaf area quantification

The extent of reporter gene expression in the *pMIR319C::GUS*, *pTCP4::GUS*, *pTCP4::TCP4-GUS*, and *pUBQ10::miR319c-GUS* transgenic lines were determined using a pixel density quantification method adapted from Kazama et al., 2010; Vercruyssen et al., 2014. A 16-bit TIFF image of the leaf primordia was imported into ImageJ (https://imagej.nih.gov/ij/). A binary version of the image was generated using the ‘make binary’ function. This function converts the coloured image into greyscale and assigns intensity values for each pixel on the grayscale 0-255 range, where zero is black, and 255 is white. Next, we used the ‘plot profile’ tool to generate a plot of the average pixel intensity (Grey value) across the width (Y-axis) for each distance from the base of the leaf primordia (X-axis). The position along the leaf length where the average pixel intensity reaches 90% of the maximum, i.e., 230, was defined as the boundary of the expression domain (denoted as the GUS domain in μm). The workflow for measuring the GUS domain is summarised with an example in **Fig. S1A**. The robustness of the pixel density based-quantification method was tested by modifying both GUS-staining times and the threshold cut-off used for determining the *pMIR319C::GUS* domain in the wildtype and the *tcp2;3;4;10* leaf primordia (**Fig. S1B, S1C**).

Areas of the dissected mature first leaves of 25-day old Col-0, *pMIR319C::MIR319C*, *mutpMIR319C::MIR319C*, and *p35S::MIR319C* T2 seedlings were determined using the polygon selection tool of the ImageJ software.

### RNA isolation and RT-qPCR

Total RNA was extracted using the TRIzol method described earlier (Challa et al., 2016). 10 μg of RNA was used for DNase (EN0521, Thermo Scientific, USA) treatment. 1.5 μg DNase-treated RNA was converted to cDNA using RevertAid Reverse Transcriptase (EP0441, Thermo Scientific, USA) and 2.5 μM oligo dT according to the manufacturer’s instructions. Quantitative PCR reactions with 25 ng cDNA and 0.4 μM primers were set up using the DyNAmo Flash SYBR Green RT-qPCR kit (F415L, Thermo Scientific, USA), according to the manufacturer’s instructions. 10 μl qPCR reactions were carried out in QuantStudio™ 6 Flex Real-Time PCR System, 384-well format (Applied Biosystems, USA). Results were analyzed using the inbuilt QuantStudio6 Pro software, and ΔΔCT values were determined after normalization with *PROTEIN PHOSPHATASE 2A SUBUNIT A3 (PP2AA3*). Relative fold changes in transcript levels were calculated using the formula 2^-ΔΔCT^. All primer sequences are provided in **Table S2**.

### Electrophoretic Mobility Shift Assay (EMSA)

EMSA was carried out as previously mentioned (Challa et al., 2016). Briefly, end-labeled oligonucleotides corresponding to class-II TCP consensus binding site were prepared with [γ-^32^P]-ATP (BRIT, India) using T4 polynucleotide kinase (EK0031, Thermo Scientific, USA). 15μL of binding reaction with 0.05 μM labeled oligonucleotide probe, 1X binding buffer (20 mM HEPES-KOH, pH 7.8, 100 mM KCl, 1 mM EDTA, 0.1% BSA, 10 ng herring sperm DNA, and 10% glycerol) and approximately 2 mg of crude recombinant protein bacterial lysate was prepared and incubated at room temperature for 30-40 min. DNA-protein complexes were analyzed on 9% native polyacrylamide gel in 0.5X TBE buffer. For super-shift assays monoclonal 0.2 μl of Anti-His antibody (H1029, Sigma-Aldrich, USA) was used. A 10 or 25-fold excess concentration of unlabelled probe was used to compete out the labeled probe. The gels were autoradiographed using a Phosphor imager (GE Typhoon FLA 9500, GE). List of oligonucleotides used for EMSA are provided in **Table S2**.

### Chromatin immunoprecipitation (ChIP) assay

ChIP was performed as previously described (Kubota et al., 2017; Challa et al., 2019). 1.5 g of fresh 10-day-old seedlings were finely grounded in liquid nitrogen. The powder was homogenized in extraction buffer-1 containing 10 mM Tris-HCl pH 8.0, 10 mM MgCl_2_, 5 mM β-mercaptoethanol, 0.1 mM PMSF, 1 mM Na_3_VO_4_, 1 mM NaF, and cOmplete protease inhibitor cocktail tablets [Roche]), followed by cross-linking with 37% formaldehyde by gentle rotation in 4°C for 10 min. 2M glycine was added to stop the cross-linking reaction. After quenching, the cross-linked samples were filtered through two layers of Miracloth (475855, Millipore, USA) and centrifuged for 10 min at 10000 g at 4 °C. The pellets containing the nuclei were resuspended and washed twice with extraction buffer-2 (0.25 M sucrose, 10 mM Tris-HCl pH 8.0, 10 mM MgCl2, 1% (w/v) Triton X-100, 5 mM β-mercaptoethanol, 0.1 mM PMSF, 1 mM Na_3_VO_4_, 1 mM NaF, and cOmplete protease inhibitor cocktail tablets) and transferred to ice-cold 1.5mL microfuge tubes. Samples were centrifuged at 14000 g for 10 min at 4 °C to isolate the nuclei. The isolated nuclei were lysed using Nuclei lysis buffer (50 mM Tris-HCl pH 8.0, 10 mM EDTA, 1% SDS, 1 mM PMSF, and cOmplete protease inhibitor cocktail tablets) and proceeded for sonication to shear the chromatin (∼ 500 bp to 1 kb size fragments) using Bioruptor Plus (Diagenode, USA) set at low mode and 45 cycles of 30 sec ON, 45 sec OFF. The sheared chromatin were diluted in ChIP dilution buffer (16.7 mM Tris-HCl pH 8.0, 167 mM NaCl, 1.1% Triton X-100, 1.2 mM EDTA, 0.1 mM PMSF, 1 mM Na_3_VO_4_, 1 mM NaF, and cOmplete protease inhibitor cocktail tablets) to make the volume up to 1.5 mL and centrifuged at 14000 g for 5 min at 4 °C. 30 μL (2%) sheared chromatin was used as input. 25 μL of Dynabeads Protein G (10004D, Invitrogen, USA) were used per sample for antibody conjugation according to the manufacturer’s instructions. For the TCP4 ChIP experiment, chromatin was immunoprecipitated using 3 μL anti-FLAG antibody (F1804, Sigma-Aldrich, USA) or Anti-IgG antibody (A9044, Sigma-Aldrich, USA) for 4 hours on a rotary shaker at 4 °C. For histone H3K27me3 ChIP experiment, chromatin was immunoprecipitated with 1 μL Anti-trimethyl-Histone H3 (Lys27) Antibody (07-449, Millipore, USA) for 16 hours on rotary shaker at 4 °C. After precipitation, the immunocomplex was washed twice with low salt buffer (20 mM Tris-HCl pH 8.0, 150 mM NaCl, 0.1% SDS, 1% Triton X-100, 2 mM EDTA), high salt buffer (20 mM Tris-HCl pH 8.0, 500 mM NaCl, 0.1% SDS, 1% Triton X-100, 2 mM EDTA) and TE buffer (10 mM Tris-HCl pH 8.0, 1 mM EDTA). Next, 50 μL of nuclei lysis buffer was added and incubated for 20 min at 65 °C to elute the immunocomplexes. Eluted samples were reverse cross-linked by adding 6 μL of 5 M NaCl and incubating at 65 °C overnight. Samples were treated with 5 μL of 10mg/ml RNase A (R6513, Sigma-Aldrich, USA) and 1 μL of 10mg/ml Proteinase K (P2308, Sigma-Aldrich, USA) and subsequently purified using QIAquick PCR purification kit (28104, Qiagen, Germany) according to manufacturer’s protocol. Extracted DNA was resuspended in 100 μL of elution buffer and proceeded for qPCR using 3μL DNA as template per reaction. Fold enrichment of TCP4 in anti-FLAG samples was expressed relative to IgG control. Fold enrichment of H3K27me3 was expressed as % Input. All ChIP qPCR primers are listed in **Table S2**.

### Yeast transformation

Yeasts were transformed with the desired DNA constructs (plasmids) by the high-efficiency LiAc-PEG method described by Walhout and Vidal, 2001, with minor modifications. A single colony of host strain was inoculated in 10 ml YPAD (Bacto yeast extract-1g, Tryptone-1g, Dextrose-2g, and Adenine sulphate-10mg per 100ml) and incubated overnight at 30 °C with constant shaking. The next day, the suitable culture volume was inoculated into 200 ml of fresh YPAD solution to an initial OD of 0.2 and kept at 30 °C with constant shaking until an OD of 0.6-0.8 was reached. Cells were pelleted down at 700 g for 10 min and washed twice with 5 ml of sterile water. The cells were washed with 2.5 ml of LiAc-TE (0.1 M LiAc, Tris-10 mM, EDTA-1mM) solution. Cells were finally resuspended in 2 ml of LiAc-TE solution. A mixture containing 280 μL of 50% PEG (Mol wt 3350, Sigma-Aldrich, USA), 36 μL of 1 M LiAc in TE, 6 μL of 10 mg/ml boiled carrier DNA (D-3159, Sigma-Aldrich, USA) was prepared and added to 1.5 ml microfuge tubes containing 60 μL cells and 100 ng of DNA construct. The cell suspension was vortexed briefly and incubated at 30 °C with constant shaking for 30 min. Cells for 30 min. Cells were transferred into a water bath maintained at 42 °C for 30 min. Cells were pelleted, washed twice with sterile water, resuspended in 100 μL sterile water, and plated onto appropriate media.

### Yeast bait, prey, and diploid generation

The full length (2.7 kb) and the non-overlapping promoter fragments, i.e., −2694-1866 (*p*^distal^), −1866-1107 (*p*^middle^), and −1107+35 (*p*^proximal^) of *MIR319C* were amplified from *pMIR319C-P4P1R* entry clone using Taq DNA polymerase and cloned into pENTR/D-TOPO vector (K240020, Invitrogen, USA) according to the manufacturer’s instructions. The entry clones were confirmed by sequencing before cloning upstream to the *minimalPro::HIS3* cassette of *pMW#2* (Gift from Ram Yadav, IISER Mohali, INDIA) by LR reaction. Y1H baits were generated according to the protocol described by Fuxman Bass et al., 2016. Briefly, the *pMIR319C* containing *pMW#2* reporter constructs were linearized with *Xho*I (R0146, New England Biolabs, USA) and integrated into the mutant *HIS3* locus of the *YM4271* host strain (Gift from Ram Yadav, IISER Mohali, INDIA) using the high-efficiency LiAc-PEG transformation method described by Walhout and Vidal, 2001. Following transformation, yeast cells were plated on the synthetic complete media lacking histidine (Sc-His) media (Walhout and Vidal, 2001) and incubated at 30 °C till colonies appeared. Post growth, genomic DNA was extracted individually from 3-4 colonies, and the integration of the *MIR319C* promoter fragment was confirmed by PCR. All primer sequences used for cloning are mentioned in **Table S2**.

For the generation of yeast diploids, each of the 384 TF-encoding plasmids was individually purified from corresponding glycerol stocks of the 1956 DEST-U Arabidopsis TF ORFeome collection (CD4-89, ABRC, USA) using QIAprep Spin Miniprep Kit. 100 ng of purified plasmid were individually transformed into *pMIR319C* baits using the high-efficiency LiAc-PEG method described earlier. Following transformation, the cells were spotted on synthetic complete media lacking histidine and tryptophan (Sc-His-Trp) media and incubated at 30 °C till colonies appeared.

### Yeast one-hybrid (Y1H) assay

The auto activity of yeast baits on Sc-His due to minimal *HIS3* expression was determined by spotting them on Sc-His media containing 0-30 mM 3-Amino-1,2,4-triazole (3-AT) (A8056, Sigma-Aldrich, USA). The minimum concentration of 3-AT that completely inhibits the growth of the bait was considered for the Y1H assay (20mM for *p^distal^*, 15mM for *p^middle^* and 15 mM for *p^proximal^*). For the co-transformation-based Y1H assay, yeast baits were transformed with TF encoding plasmids and selected on Sc-Trp-His media. Following selection, the cells were resuspended in Sc-Trp-His media and allowed to grow in shaking incubators for 24 hours at 30 °C. After 24 hours of growth, OD was measured. Volumes corresponding to equal OD were spotted on Sc-Trp-His media with respective concentrations of 3-AT and incubated at 30 °C. The plates were inspected for growth and imaged over days 3-5 post-incubation.

### Statistical analysis

All graphical representations and statistical analysis were performed using GraphPad Prism 9 software (GraphPad, USA). First, samples were tested for Gaussian distribution using the Shapiro-Wilk normality test. If samples were normally distributed, pairwise comparisons were performed using unpaired *t-*test. Welch’s correction was applied if variances were different among the samples. Statistical differences among multiple genotypes were determined by ordinary One-Way ANOVA followed by Tukey’s multiple comparisons test or One-Way ANOVA followed by Dunnet’s T3 post-hoc test, depending on whether variances were significantly different among the genotypes. If the samples failed the normality test, then pairwise comparisons were carried out by Mann Whitney test. Statistical differences among multiple genotypes were determined by One-Way ANOVA followed by Dunn’s post-hoc test. To compare statistical differences among multiple genotypes subjected to ethanol (Mock) or dexamethasone (DEX) treatments, we used Two-Way ANOVA followed by Tukey’s post-hoc test. *p-*values obtained from the respective tests are indicated above the data bars in majority of the plots. For plots with a large number of comparisons, statistical differences among samples are indicated by different alphabets.

### Accession numbers

TAIR (The Arabidopsis Information Resource, www.arabidopsis.org) accession numbers of the major genes used in this study are *AT4G23713* (*MIR319A/JAW*), *AT2G40805 (MIR319C), AT4G18390 (TCP2), AT1G53230 (TCP3), AT3G15030 (TCP4), AT2G31070 (TCP10), AT1G30210 (TCP24), AT3G11940 (RPS5A), AT4G05320 (UBQ10)*, *AT1G13710 (CYP78A5/KLUH), AT3G59900 (ARGOS)*, *AT1G13320* (*PP2AA3*), *BLS (AT3G49950)*, and *AT3G45140 (LOX2)*.

## Acknowledgements

We thank Thomas Jack (Dartmouth College, Hanover, USA) for *pMIR319C::GUS* line, Tsuyoshi Nakagawa (Shimane University, Japan) for gateway DNA constructs for generating *GUS*-based reporter lines, Ram Yadav (IISER Mohali, India) for yeast strains and gateway DNA constructs for Y1H assay. Anurag N Sharma (IISc, Bengaluru, India) is acknowledged for generating the *tcp2;3;4;10* line.

## Author contributions

N.S. initiated the project and performed almost all the experiments, analyzed and interpreted the results, organized the figures, wrote the first draft of the manuscript and contributed to its finalization; P.S. generated the Col-0;*GR^#2^* and the Col-0;*pTCP3::rTCP3-GR* transgenic lines that have been very useful for this study and helped in drawing the primary conclusion; U.N. contributed to designing experiments and data interpretation, guided the first two authors, corrected the manuscript and finalized it.

## Supplementary figures

**Figure S1.**
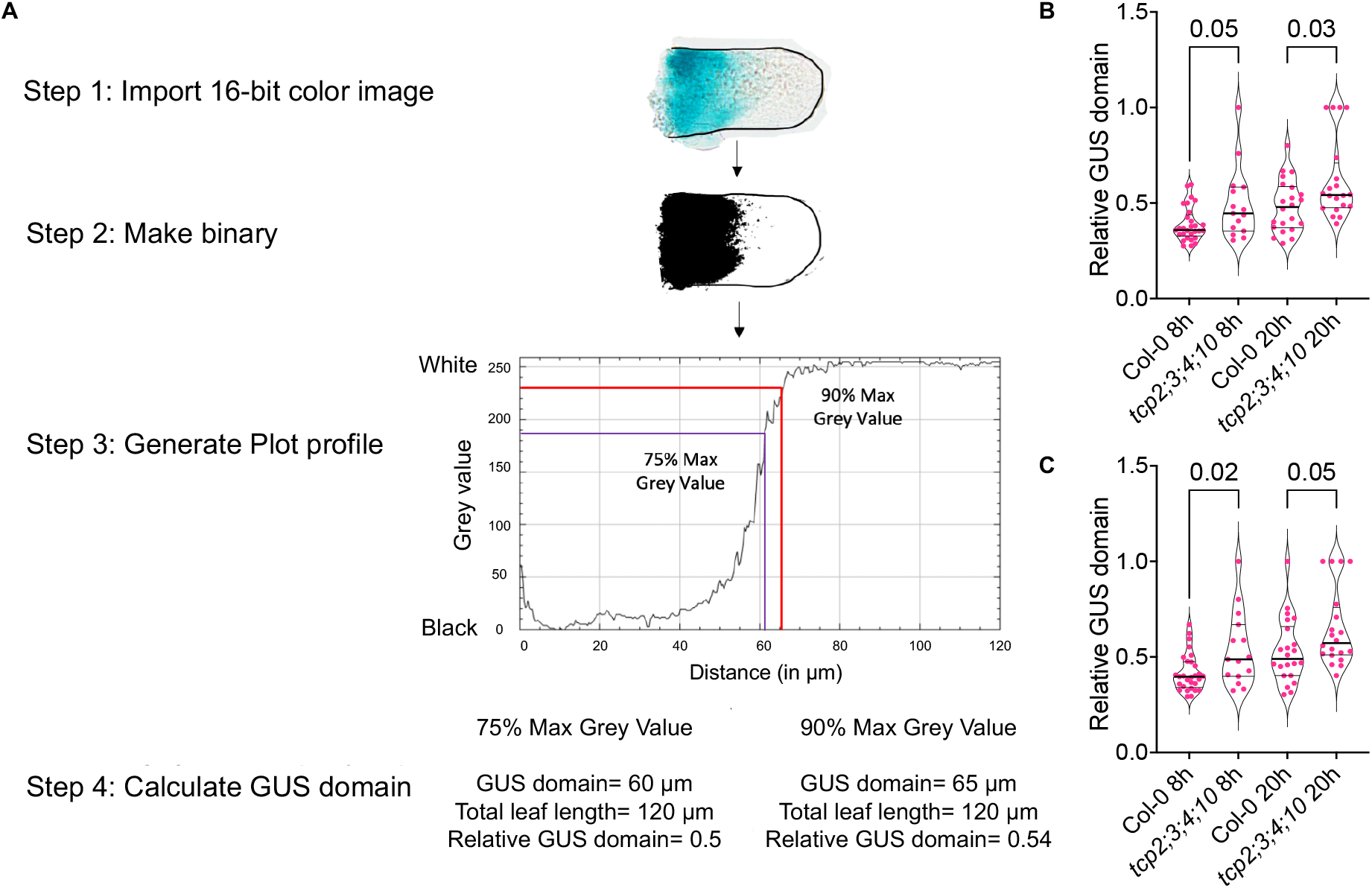
(Supports Figures 1 and 2): Method for quantifying *GUS* expression domains in leaf primordia. **(A)** Schematic representation of the pixel density-based method for quantification of *GUS* expression domain in leaf primordia. Domain of *GUS* activity was determined using the plot profile function of the ImageJ software (https://imagej.nih.gov/nih-image/more-docs/Tutorial/Profile.html). **(B-C)** Violin plots displaying the proportion of *pMIR319C::GUS* domain relative to leaf length in Col-0 and in *tcp2;3;4;10* leaf primordia of length 50-350 µm. GUS-staining was performed either for 8 hours or for 20 hours, as indicated on the X-axis labels. The proportion of GUS domain determined using 75% **(B)** or 90% **(C)** intensity cut off. N=15-29 leaves. Differences among samples are indicated by *p*-values on top of the comparisons, as determined by Mann-Whitney test.

**Figure S2.**
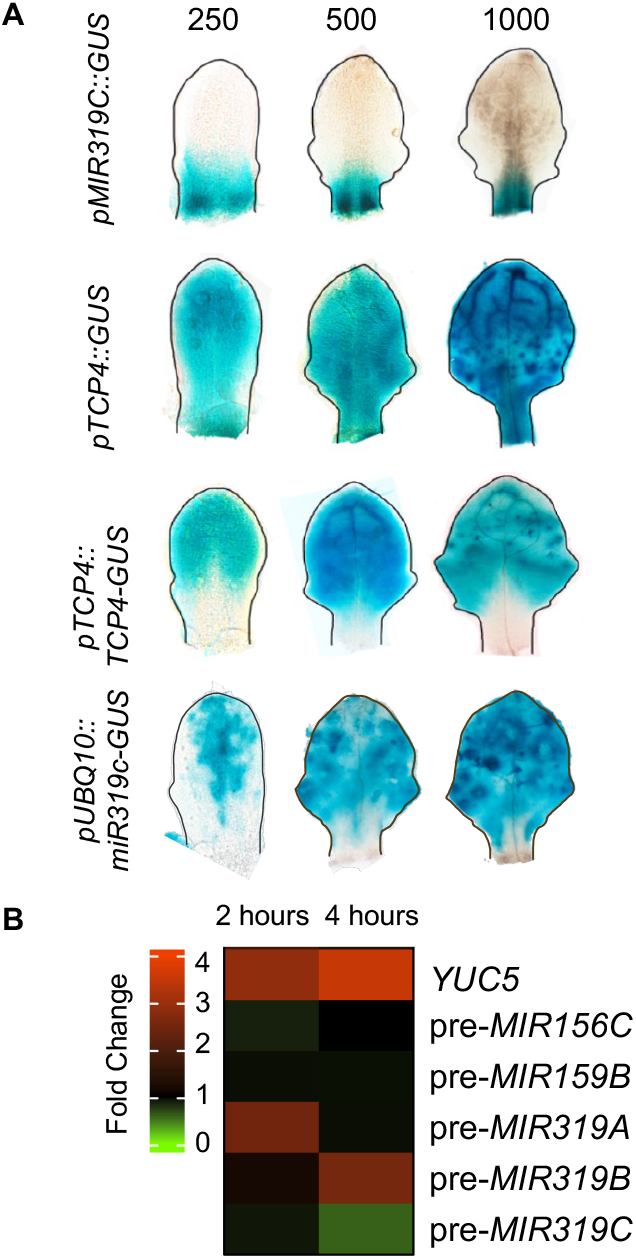
(Supports Figure 1): Expression pattern of *MIR319C* and *TCP4* in young leaves. **(A)** Bright field images of 250, 500 and 1000 μm long wild-type leaf primordia expressing reporter constructs for *MIR319C* promoter (*pMIR319C::GUS*), *TCP4* promoter (*pTCP4::GUS*), TCP4 protein (*pTCP4::TCP4-GUS*), and miR319 activity (*pUBQ10::miR319c-GUS*) respectively. (**B)** Heatmap displaying change in transcript levels of indicated genes in 9-day old *jaw-D;pTCP4::rTCP4-GR* (*jaw-D;GR^#1^*) (Challa et al., 2016) seedlings treated with 12 µM dexamethasone (DEX) for 2 or 4 hours. pre-*MIR319C* level is specifically downregulated upon 4 hours of DEX treatment as opposed to pre-*MIR319A* and pre-*MIR319B* (both upregulated) or pre-*MIR156C* and pre-*MIR159B* (unchanged). *YUC5*, a direct target of TCP4 (Challa et al., 2016), is used as a positive control. This is a reanalysis of microarray data reported in Challa et al., 2016.

**Figure S3.**
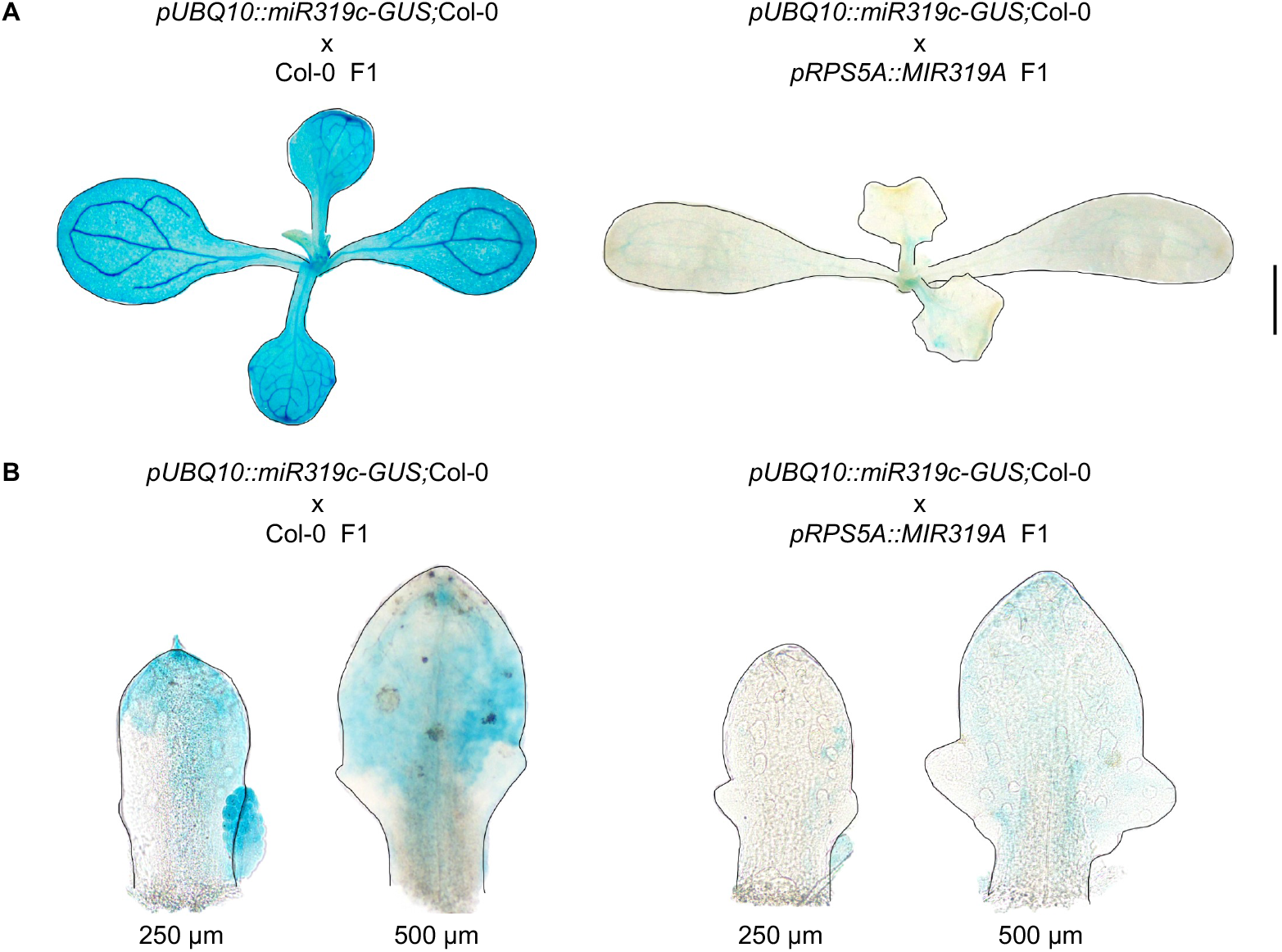
(Supports Figure 1): *pUBQ10::miR319c-GUS* domain reflects miR319c activity. **(A-B)** Bright field images of **(A)** 9-day old whole seedlings and **(B)** 250 µm and 500 µm long leaf primordia at the 3^rd^ and 4^th^ nodes of 9-day old Col-0 and *pRPS5A::MIR319A* heterozygote individuals expressing *pUBQ10::miR319c-GUS* reporter. Scale bar in **(A),** 1 mm.

**Figure S4.**
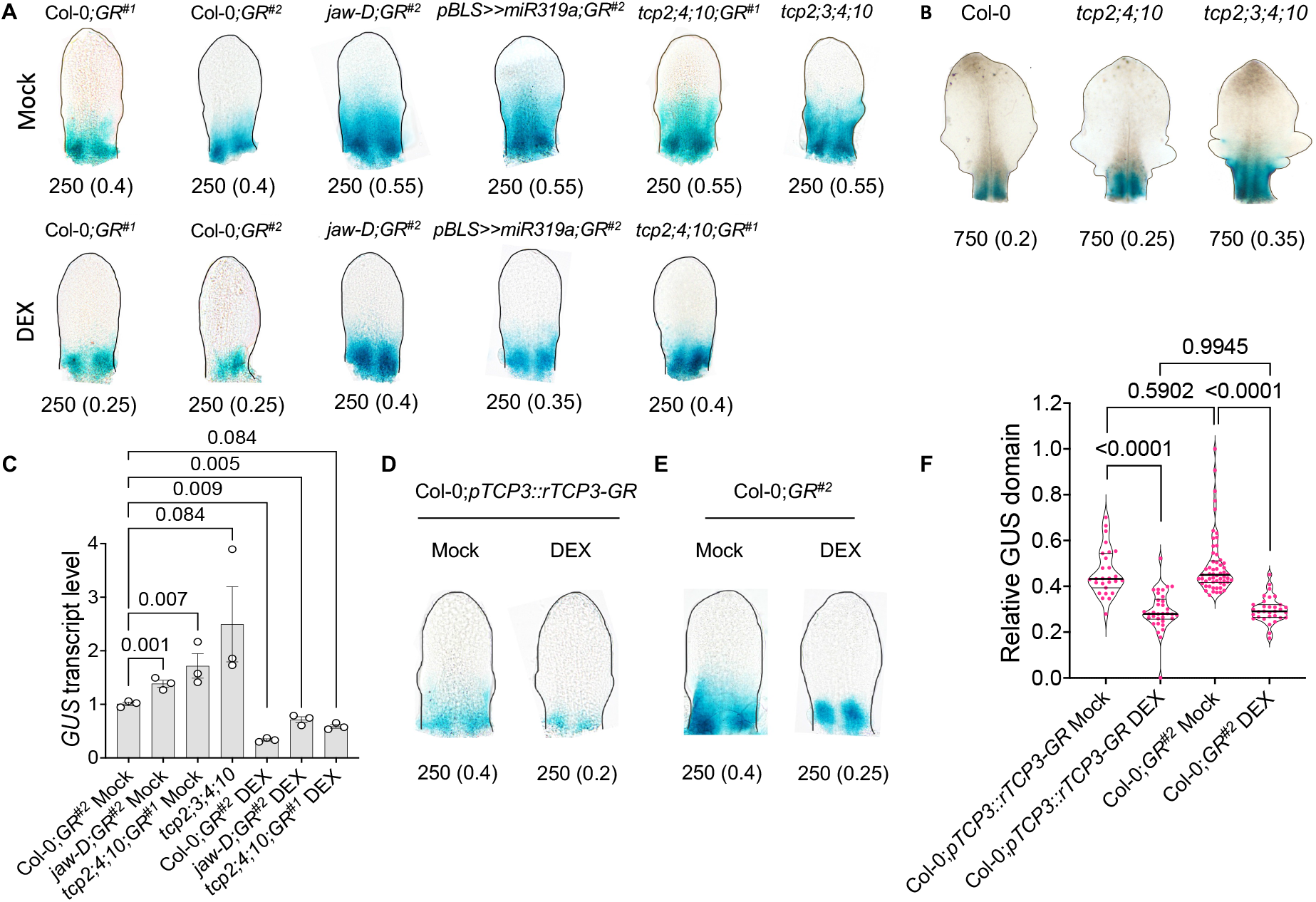
(Supports Figure 2): Altered *MIR319C* promoter activity upon JAW-TCP manipulation in the distal region of early leaf primordia. **(A)** Bright field images of the ethanol (Mock) or 6 µM dexamethasone (DEX)-treated 6^th^ leaf primordia of the indicated genotypes expressing *pMIR319C::GUS* transgene. Numbers below the images indicate leaf length in µm and GUS domain/leaf length (in parentheses), respectively. **(B)** Bright field images of 750 µm long leaf primordia at the 3^rd^ nodes of seedlings of indicated genotypes expressing *pMIR319C::GUS* reporter. Numbers below the images indicate leaf length in µm and GUS domain/leaf length (in parentheses), respectively. (C) *GUS* transcript level in the 9-day old whole seedlings of indicated genotypes treated to 0 µM (Mock) or 6 µM (DEX) dexamethasone. N=3 biological replicates, each containing ∼100 mg of tissue. Error bars indicate SEM. Differences among samples are indicated by *p*-values on top of the comparisons, One-way ANOVA, Dunnett’s post hoc test was performed. **(D-E)** Bright field images of 6^th^ leaf primordia of the indicated genotypes expressing *pMIR319C::GUS* transgene. Mock and DEX in **(D)** and **(E)** indicate treatment with ethanol or 6 µM dexamethasone, respectively. The numbers below the images indicate leaf length. Proportion of GUS domain relative to leaf length is shown in the parentheses. **(F)** Violin plot representations of the relative *pMIR319C::GUS* domain (N=27-57) in 50-250 µm long primordia on 7^th^-8^th^ nodes of seedlings of indicated genotypes treated without (Mock) or with (DEX) 6 µM dexamethasone. Differences among samples are indicated by *p*-values on top of the plots. Two-way ANOVA, Tukey’s post hoc test was performed.

**Figure S5.**
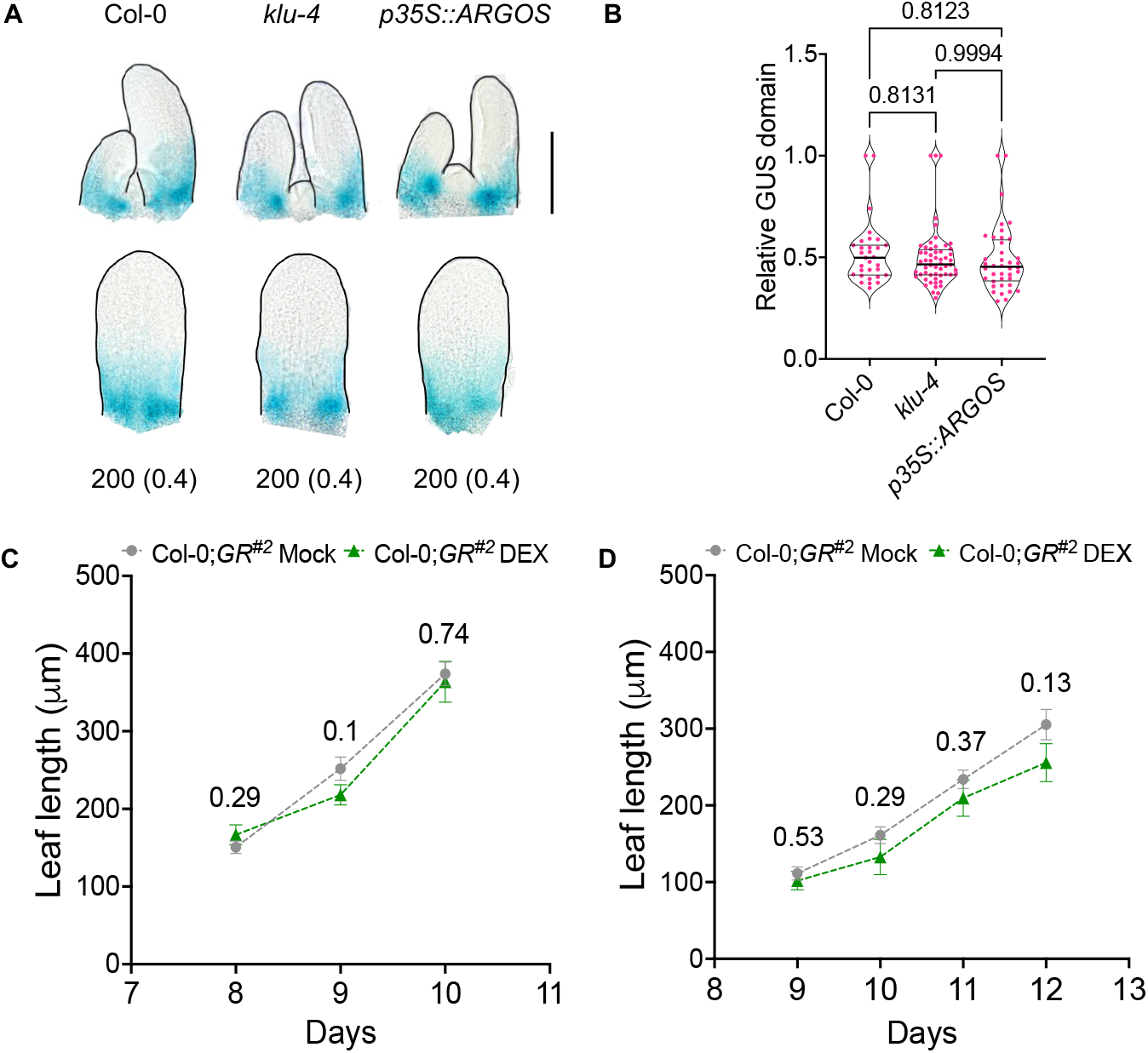
(Supports Figure 2): *MIR319C* expression is independent of cell proliferation in early leaf primordia. **(A)** Bright field images of 10-day old shoot apices and 6^th^ leaf primordia of the indicated genotypes expressing *pMIR319C::GUS* transgene. Scale bar for the top panel, 100 μm. Numbers below the images refer to leaf length in µm. Proportions of GUS domain relative to leaf length are shown in the parentheses. **(B)** Violin plots displaying the distribution of *pMIR319C::GUS* domain/leaf length in leaf primordia of the genotypes indicated on the X-axis. N=31-56 leaves. Differences among samples are indicated by *p*-values on top of the comparisons, One-way ANOVA, Dunnett’s post hoc test was performed. **(C) and (D)** Scatter plots showing mean lengths of the ethanol-treated (Mock) or 6 µM dexamethasone (DEX)-treated Col-0;*GR^#2^* leaf primordia on corresponding days after stratification. Each grey circle or green triangle represents the mean lengths of 6^th^ leaf primordia on 8-10 days **(C)** and 8^th^ leaf primordia on 9-12 days **(D)** after stratification (indicated on X-axis). N=10-35 **(C)** and 8-25 **(D)** leaves. Error bars indicate SEM. Numbers above the error bars correspond to the *p*-values derived from unpaired *t*-test with Welch’s correction for differences in variance.

**Figure S6.**
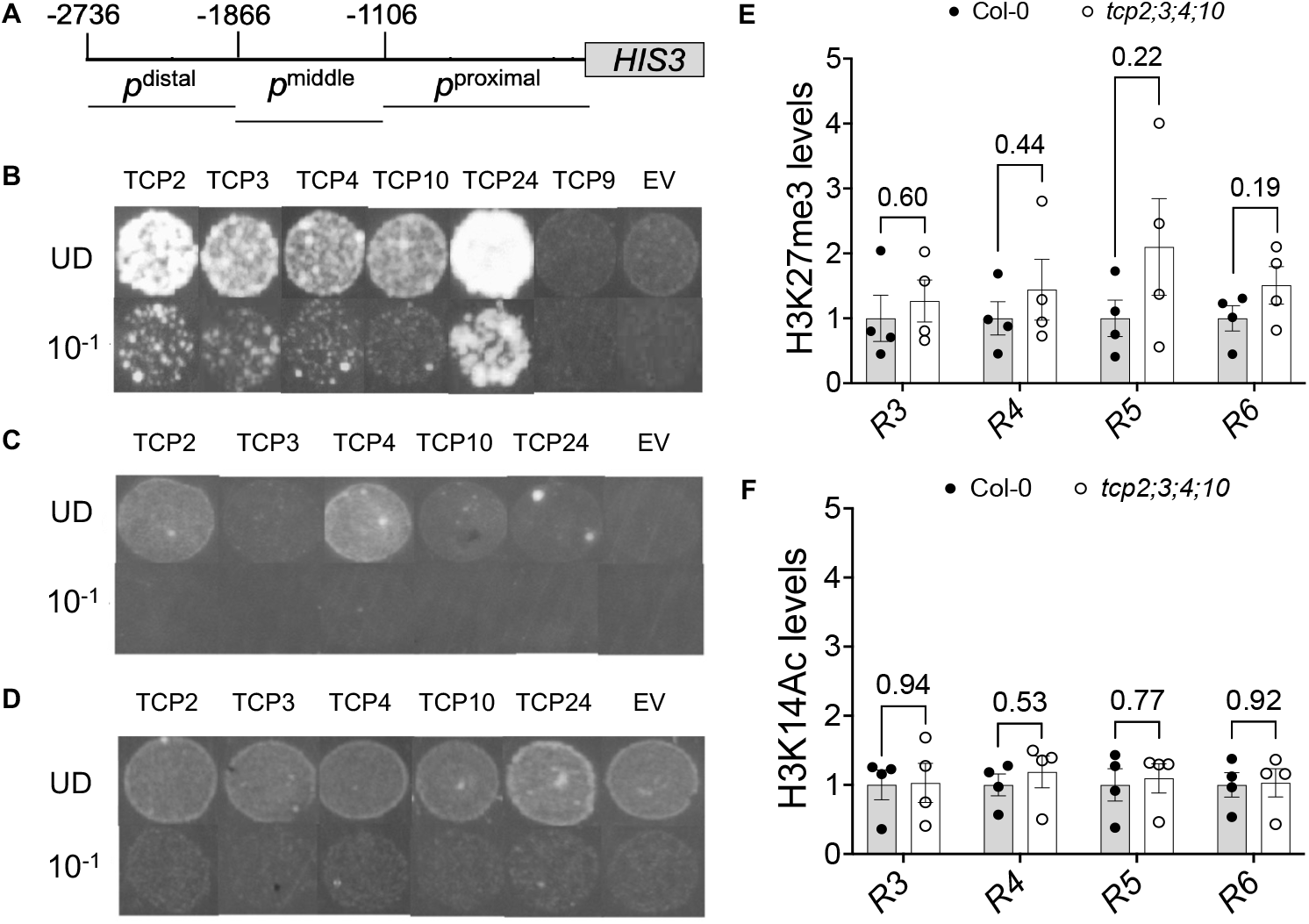
(Supports Figure 4): TCP4 binds specifically to the distal URR element of *MIR319C* locus. **(A)** Schematic representation of the *HISTIDINE3* (*HIS3*)-based reporter construct containing 2736 bp *MIR319C* URR used for yeast bait generation. Grey box indicates *HIS3 CDS* sequence. The three horizontal lines below the URR scheme represent the truncated promoter regions, i.e., *p*^distal^ (−2736 to −1866), *p*^middle^ (−1866 to −1106) and *p*^proximal^ (−1106) used to generate the bait constructs for the yeast one-hybrid assay. **(B-D)** Images of yeasts harboring three truncated *MIR319C* promoter regions, *p*^distal^ **(B)**, *p*^middle^ **(C)** or *p*^proximal^ **(D)** upstream to the *HIS3* reporter gene and corresponding to preys as indicated, grown in the presence of inhibitory concentrations (15-20 µM) of the *HIS3* inhibitor 3-amino-1,2,4-triazole (3-AT). Yeast culture suspensions (OD_600_ value 0.3) were spotted on Sc-His-Trp+3-AT media undiluted (UD) and at a dilution of 10^-1^. EV, empty vector negative control. **(E) and (F)** Fold enrichment of the fragments corresponding to R3-R6 regions precipitated by anti-H3K27me3 **(E)** and anti-H3K14Ac **(F)** antibody in the ChIP experiment with fragments precipitated from chromatin preparations of 9-day old Col-0 and *tcp2;3;4;10* seedlings. Fold change in H3K27me3 **(E)** and H3K14Ac **(F)** levels represented as % Input and normalized to values from respective Col-0 controls. Error bars represent SEM of four biological replicates (performed in technical duplicates). Pairwise comparisons were performed using unpaired *t*-test. *p*-values are shown above the data bars.

**Figure S7.**
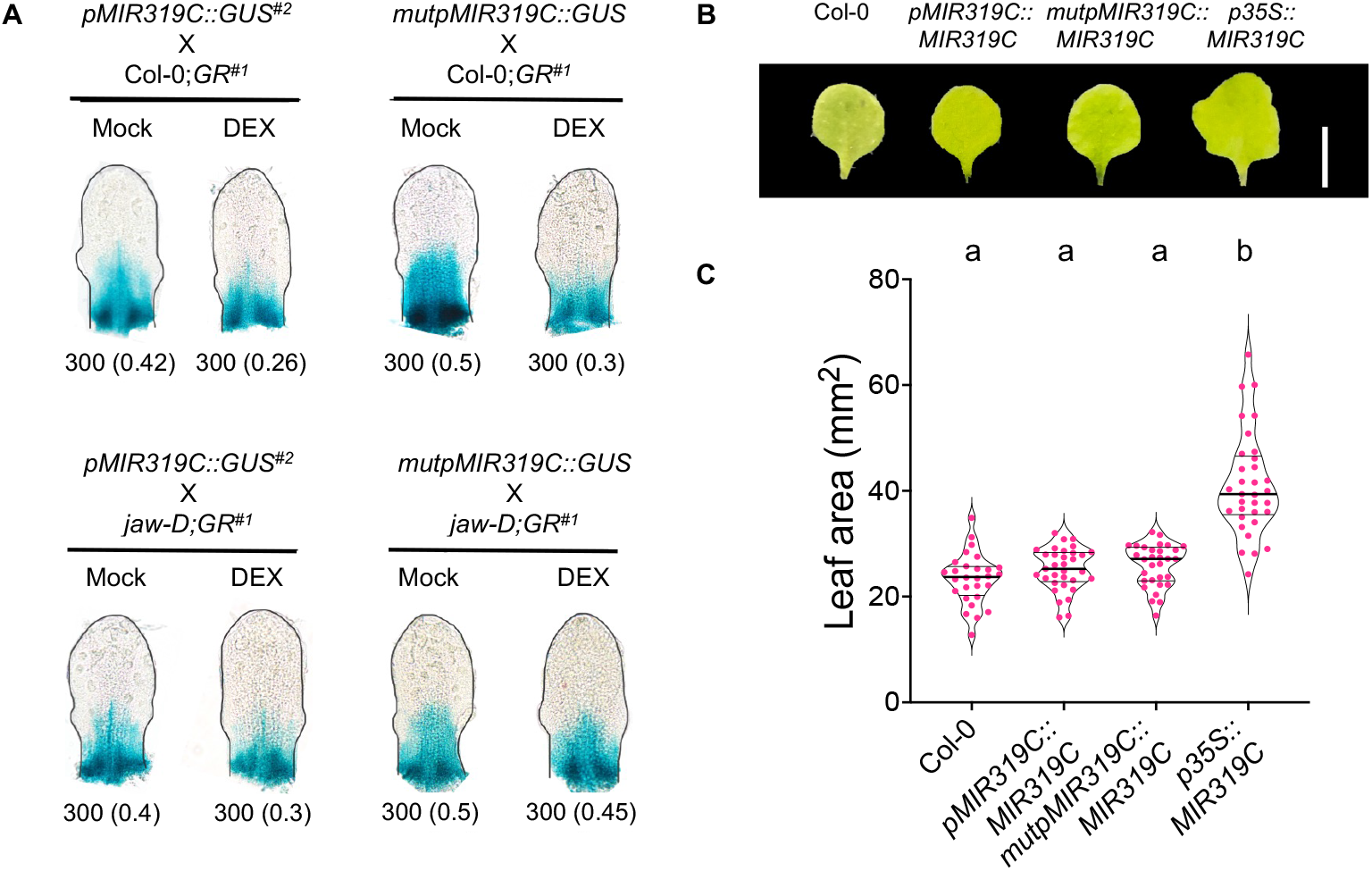
(Supports Figure 5): Effect of mutating the TCP4-binding elements on *MIR319C* promoter activity and mature miR319 activity. **(A)** Bright field images of the 5^th^ leaf primordia from Col-0;*GR^#1^* (top panel) and *jaw-D;GR^#1^* (bottom panel) seedlings expressing *pMIR319C::GUS^#2^*or *mutpMIR319C::GUS* transgenes as indicated. Numbers below the leaf primordia images indicate leaf length and GUS domain/leaf length (in parentheses), respectively. **(B)** Representative images of the 1^st^ leaf from 25-day old seedlings of the indicated genotypes. Scale bar, 5 mm. **(C)** Area of the 1^st^ leaf of the genotypes indicated on the X-axis. N=28-33 leaves. Differences among samples are indicated by different alphabets on top of the violin plots, p<0.001. One-way ANOVA, Tukey’s post hoc test was conducted.

## Supplementary Tables

**Table S1:** Genotypes used for this study.

| Genotype | Description | Source |
| --- | --- | --- |
| Columbia-0 (Col-0) | Wild type | NA |
| *jaw-D* | Overexpression line for miR319a under CaMV 35S enhancer | Palatnik et al.,2003 |
| *pBLS>>miR319a* | Overexpression line for miR319a under *BLS* promoter | Efroni et al., 2008 |
| *tcp2;4;10* | Knockout line for *TCP2*, *TCP4*, and *TCP10* | Challa et al., 2019 |
| *tcp2;3;4;10* | Knockout line for *TCP2*, *TCP3*, *TCP4*, and *TCP10* | This study |
| Col-0;*pTCP4::rTCP4-GR<sup>#1</sup>*<br>(Col-0;*GR<sup>#1</sup>*) | Dexamethasone Inducible line#1 for *rTCP4* activity under *TCP4* promoter | Challa et al., 2016 |
| *klu-4* | Knock out line for *CYP78A5/KLUH* | Anastasiou et al., 2007 |
| *p35S::ARGOS* | Overexpression line for *ARGOS* under CaMV 35S promoter | Hu et al., 2003 |
| *p35S::TCP4-3F6H* | Overexpression of 3XFLAG-6XHis tagged *TCP4* under CaMV 35S promoter | Kubota et al., 2017 |
| *pMIR319C::GUS<sup>#1</sup>* | GUS reporter line#1 for *MIR319C* promoter activity | Nag et al., 2009 |
| *pTCP4::GUS* | GUS reporter line for *TCP4* promoter activity | Challa et al., 2019 |
| *pTCP4::TCP4-GUS* | GUS reporter line for *TCP4* protein accumulation | Challa et al., 2019 |
| Col-0;*pTCP4::rTCP4-GR<sup>#2</sup>*<br>(Col-0;*GR<sup>#2</sup>*) | Dexamethasone Inducible line#2 for *rTCP4* activity under *TCP4* promoter | This study |
| Col-0;*pTCP3::rTCP3-GR* | Dexamethasone Inducible line for *rTCP3* activity under *TCP3* promoter | This study |
| *pMIR319C::rTCP4-GR<sup>#1.1;#1.2</sup>* | Dexamethasone Inducible line#1 and #2 for *rTCP4* activity under *MIR319C* promoter | This study |
| *pRPS5A::rTCP4-GR<sup>#1.1;#1.2</sup>* | Dexamethasone Inducible line#1 and #2 for *rTCP4* activity under *RPS5A* promoter | This study |
| *pMIR319C::MIR319C* | Overexpression of miR319c under *MIR319C* promoter | This study |
| *mutpMIR319C::MIR319C* | Overexpression of miR319c under *MIR319C* promoter carrying mutations in JAW-TCP binding sites | This study |
| *p35S::MIR319C* | Overexpression of miR319c under CaMV 35S promoter | This study |
| *pRPS5A::MIR319A* | Overexpression of miR319a under *RPS5A* promoter | This study |
| *pMIR319C::GUS<sup>#2</sup>* | GUS reporter line#2 for *MIR319C* promoter activity | This study |
| *mutpMIR319C::GUS*, | GUS reporter line for activity of *MIR319C* promoter carrying mutations in JAW-TCP binding sites | This study |
| *pUBQ10::miR319c-GUS* | GUS reporter line for miR319c activity | This study |

**Table S2:**
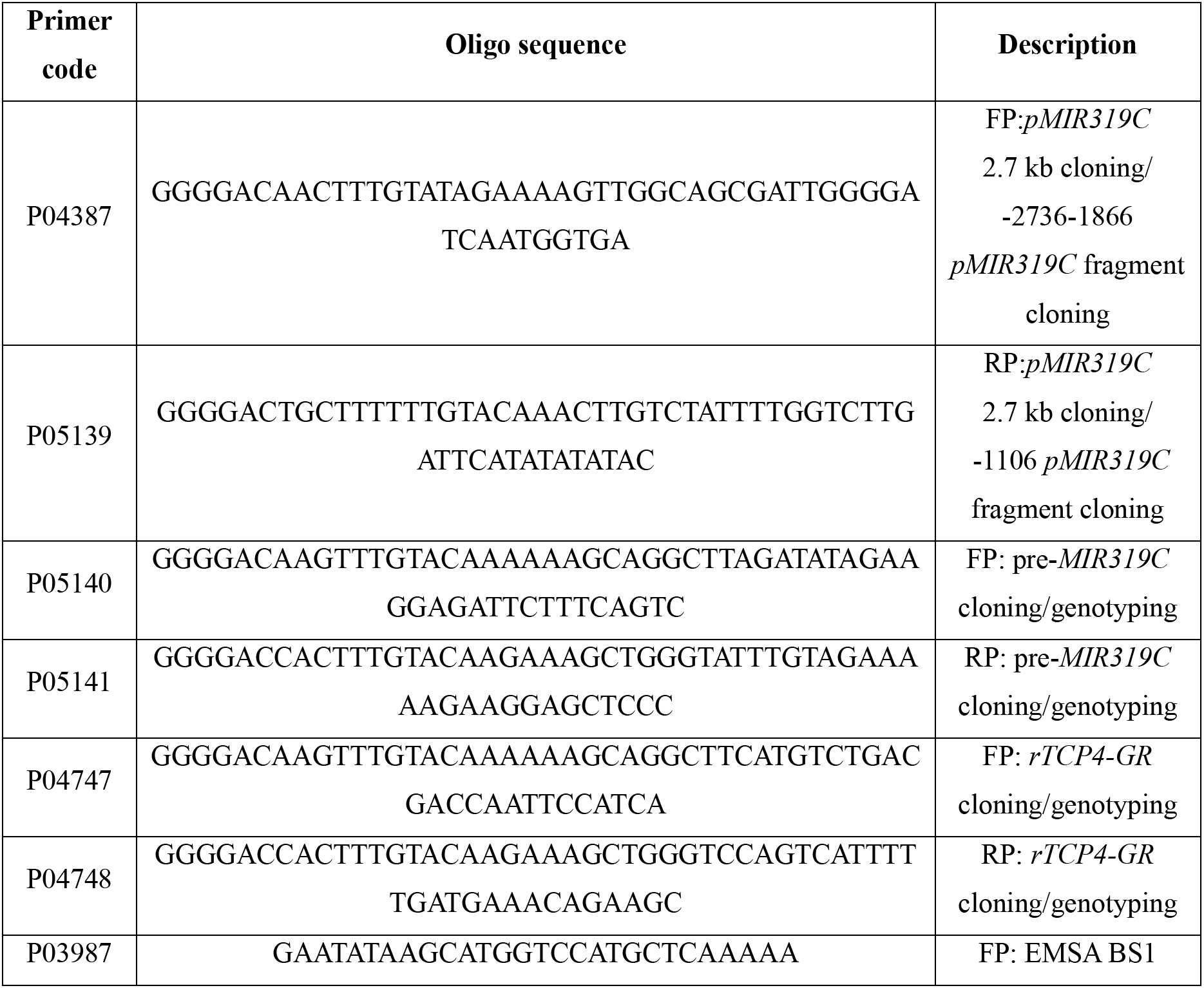
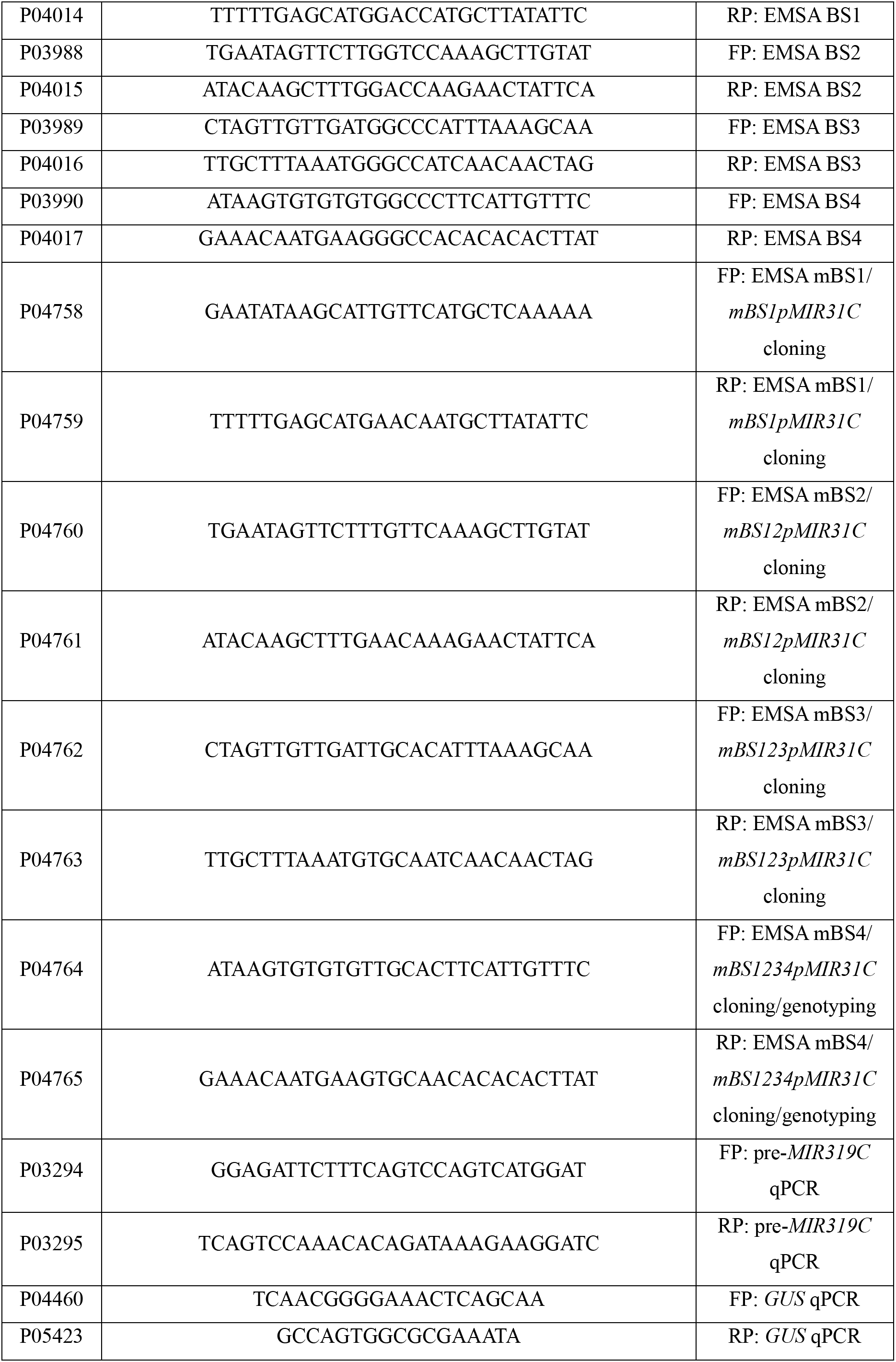
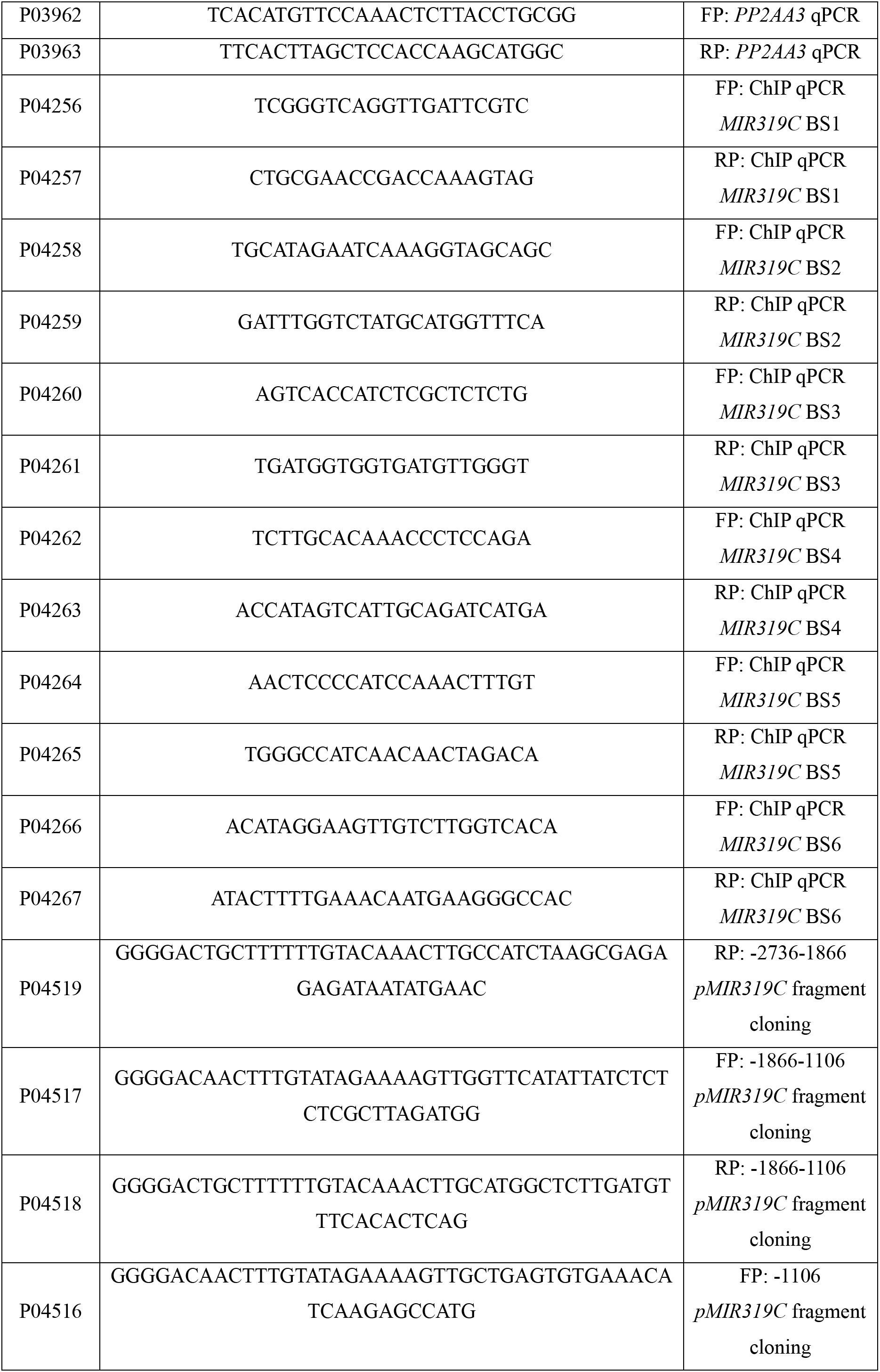
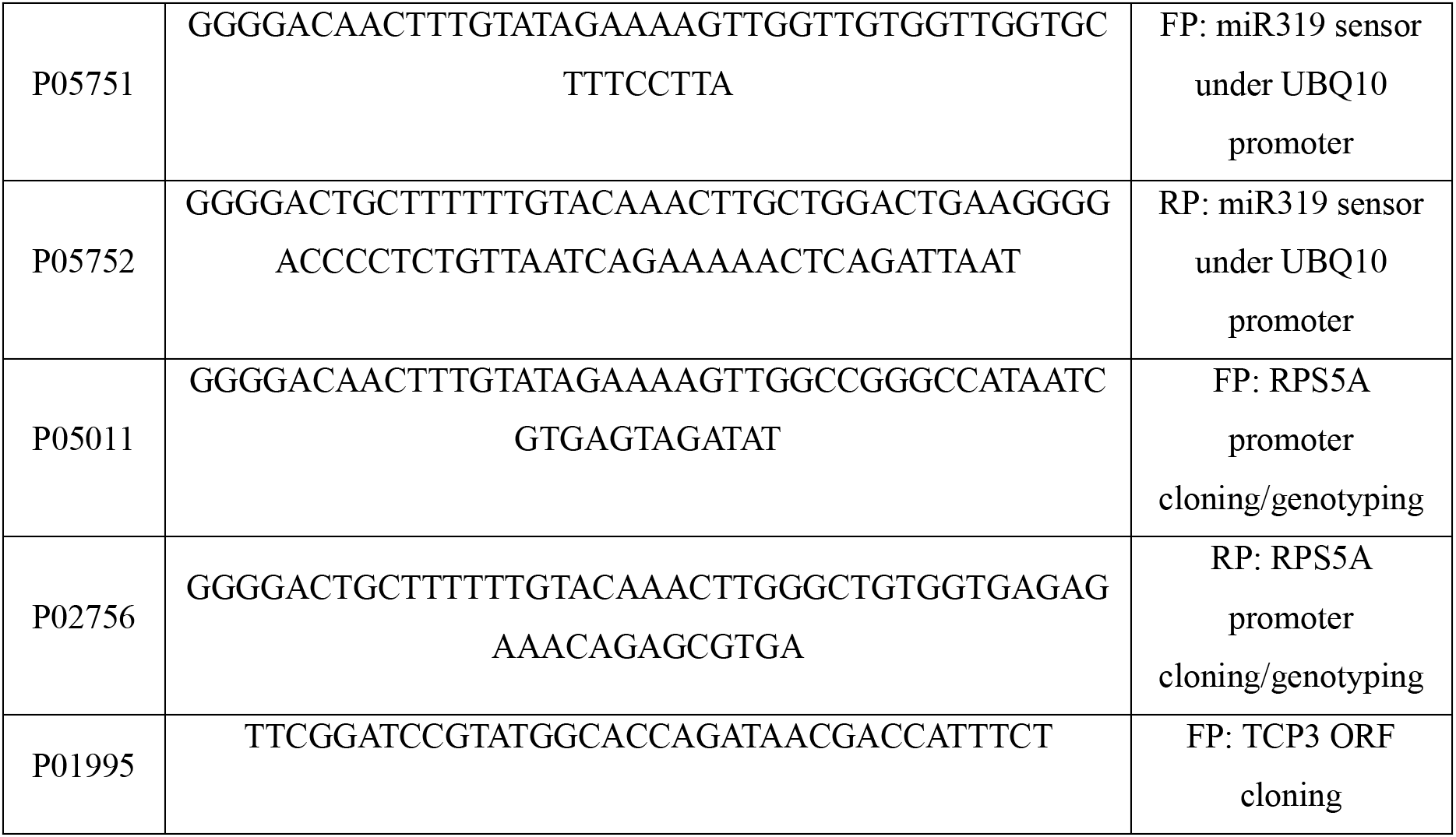
Oligonucleotides used for this study (Oligos are written in 5’ 3’ direction. FP and RP refer to forward primer and reverse primer, respectively)

